# Syntaxin of plants 32 regulates pollen wall development and pollen tube cell wall integrity via controlling secretory pathway

**DOI:** 10.1101/2023.02.03.527076

**Authors:** Yuqi Liu, Xiaonan Zhao, Guangtao Qian, Xiaohui Ma, Minglei Song, Guochen Qin, Shanwen Sun, Mingyu Wang, Kaiying Gu, Wei Sun, Jian-Kang Zhu, Lixi Jiang, Lixin Li

## Abstract

Pollen tubes (PTs) elongate in a polar way to deliver sperm cells to the ovule. Pollen wall development and PT cell wall integrity (CWI) maintenance are critical for PT growth and double fertilization. Pollen wall development mainly relies on secretion of exine precursors in tapetum. RALF4/19-ANX/BUPS-MRI and RALF4/19-LRX-AUN are two distinct signaling pathways but converge to fine-tune CWI during PT growth. Here, we discovered that *atsyp32+/-*, *AtSYP32* RNAi and *AtSYP3132* RNAi lines were male sterile. The tapetum development in these lines were disturbed, and the pollen wall structure was impaired resulting in pollen grain and tube bursting and less PTs navigated to micropyles. Strikingly, there were numerous ectopic secretory vesicles retained in pollen cytoplasm, and the abundance or distribution of polysaccharides and AGPs altered significantly in PTs of the mutants and RNAi lines. AtSYP32 interacted with the vesicle transport regulators SEC31B, SEC22 and BET12, the PT CWI regulators RALF19 and LRX11, and the XyG xylosyltransferase XXT5, in the Golgi apparatus. Transcription of some genes related to pollen wall biosynthesis and PT CWI maintenance were seriously affected by *AtSYP32* downregulation. Our findings illustrate that AtSYP32 plays essential roles in pollen wall development and PT CWI maintenance via controlling secretory pathway.

**IN A NUTSHELL:** *Background:* Pollen wall is the most complex cell wall. Pollen wall development mainly relies on secretion of precursors of exine and pollen coat in tapetal cells. Pollen tubes (PTs) grow in a polar way to deliver sperm cells to the ovule. Maintenance of PT cell wall integrity (CWI) is critical for PT elongation and double fertilization. RALF4/19 ligands interact with BUPS-ANX receptors, signaling it in an autocrine manner to maintain CWI during PT elongation. RALF4/19-LRX-AUN pathway is distinct with RALF4/19-ANX/BUPS-MRI pathway but they converge to fine-tune CWI during PT growth. Biosynthesis of PT cell wall involves multiple subcellular compartments and vesicle transport pathways. Golgi apparatus acts as a hub in vesicle trafficking. Golgi-syntaxin AtSYP31 and AtSYP32 regulate pollen development by controlling intra-Golgi transport and Golgi morphology

*Question:* What is AtSYP32 role in pollen wall and tapetum development? Who are the AtSYP32 partners that regulate secretion of cell wall biosynthesis materials?

*Findings:* We found that no homozygote progeny was obtained from self-pollinated *atsyp32+/-* alleles due to pollen sterile. The tapetum development and degeneration in *atsyp32+/-* mutants was severely delayed, and the pollen wall and PT wall structure were impaired. Strikingly, there were numerous ectopic secretory vesicles retained in pollen cytoplasm in *atsyp32+/-* mutants, and the abundance or distribution of PT wall polysaccharides and AGPs altered obviously. AtSYP32 interacted with the vesicle transport regulators SEC31B, SEC22 and BET12, the PT CWI regulators RALF19 and LRX11, and XyG xylosyltransferase XXT5, in the Golgi. All these highlight that AtSYP32 regulates pollen wall development and maintenance of PT CWI via controlling secretory pathway.

*Next steps:* The biological significances and the molecular mechanisms of AtSYP32 interacting with XXT5, RALF19 and LRX11 are elusive but thought-provoking. We are going to clarify the mechanisms.

## Introduction

Gametophyte development is crucial for sexual reproduction in higher plants. Pollen tubes (PTs) elongate in a polar way to deliver sperm cells to the ovule for double fertilization (Johnson et al., 2019). The elongation mechanism requires tight coordination between the cell wall remodeling, cytoskeleton dynamics, secretory pathways and signal transduction (sumarized by Guo and Yang, 2020). The PT cell wall needs to be robust to withstand turgor pressure and protect cell structure and functions. It is highly dynamic to support cell growth and transduce intracellular and extracellular signals (Hepler et al., 2013). The PT cell wall is composed of callose, nonesterified pectins, glycoproteins, and cellulose-like polymers (Chebli et al., 2012). At the PT growing tips, cell wall remodeling is tightly controlled by coordination of the glycoproteins and structural materials in the apoplast (Hepler et al., 2013).

The maintenance of cell wall integrity (CWI) of PT is critical for double fertilization. The mechanism is a complicated signaling dialogue which is mediated by the *Catharanthus roseus* receptor like kinase 1-like (CrRLK1L) receptor family and the rapid alkalinization factor (RALF) ligand peptides (Ge et al., 2017, 2019). CrRLK1Ls are transmembrane receptors that sense external signals and trigger intracellular signaling to direct cell growth. Therefore, The CrRLK1L receptors have been established as key regulators of CWI (Nissen et al., 2016; Franck et al., 2018). CrRLK1L family receptors ANXUR1 (ANX1), ANX2, Buddha’s paper seal1 (BUPS1) and BUPS2 are essential for PT CWI. *anx1 anx2* double mutant exhibits precocious PT burst leading to male sterile (Miyazaki et al., 2009; Ge et al., 2017, 2019). *bups1 bups2* double mutant is deficient on male transmission (Ge et al., 2017). ANX1/2 interact with BUPS1/2 to form a heteromeric receptor complex to maintain PT CWI (Ge et al., 2017; Ge et al., 2019). MARIS (MRI), a receptor-like cytoplasmic kinase (RLCK), is involved in maintenance of PT CWI. *mri* mutant PTs ruptured prematurely and results in pollen sterile (Boisson-Dernier et al., 2015). MRI functions link to ANX-mediated signaling pathway since *MRI (R240C)* but not wild type *MRI* expression can partially rescue PT bursting phenotype in *anx1 anx2* (Boisson-Dernier et al., 2015).

The RALF peptides have been demonstrated as ligands for the CrRLK1L family receptors (Ge et al., 2017). RALF4 and 19 function redundantly in maintenance of PT CWI. *ralf4 ralf19* double mutant displays precocious PT burst resembling *anx1 anx2* and *bups1 bups2* phenotypes. RALF4 and 19 interact with the BUPS-ANX complex, signaling it in an autocrine manner to maintain CWI during PT elongation in the pistil (Ge et al., 2017; Mecchia et al., 2017). Leucine-rich repeat (LRR) extensin (LRX), a cell wall glycoprotein, contains a C-terminal extensin domain and an N-terminal LRR domain. The extensin domain anchors LRXs to the cell wall, and LRR domain seems to associate with the plasma membrane (PM) and potentially transmit signals by binding partner proteins with signaling capacity (Wang et al., 2018). In *Arabidopsis*, LRX family proteins are divided into two classes, LRX1-7 belong to vegetative class (Zhao et al., 2021); and LRX8-11 belong to reproductive class which is specifically expressed in pollen and PTs (Wang et al., 2018). The triple mutants and a quadruple mutant of *LRX8–11* show significantly reduced male transmission because of burst pollen grains and tubes, indicating the essential roles of LRX8–11 in PT growth and CWI maintenance (Wang et al., 2018). RALF4 binds LRXs and negatively regulates PT growth. *lrx8-11* triple and quadruple mutants are insensitive to RALF4-induced inhibition of PT growth (Mecchia et al., 2017). ATUNIS1 (AUN1) and AUN2, the protein phosphatases, are negative regulators of PT growth. AUN1 functions downstream of LRX8-11, a pathway parallel to that mediated by ANX1/2–MRI. The two distinct pathways converge to fine-tune CWI during tip growth (Franck et al., 2018). The female-derived ligand RALF34 competes with RALF4/19 to bind BUPS-ANX receptor complex at the interface of PT-female gametophyte contact site to induce PT bursting and sperm release (Ge et al., 2017). FERONIA (FER), a CrRLK1L receptor kinase expressed in the female gametophyte, is essential for PT reception (Stegmann et al., 2017). When PT enters micropyle, hercules receptor kinase 1 (HERK1)/ANJ-FER-LRE complex in the synergid cells is activated by an unknown ligand and improves contents of Ca^2+^ and ROS to induce PT bursting (Somoza et al., 2021).

In *Arabidopsis*, the pollen wall consists of intine, exine, and tryphine (pollen coat) (Gómez et al., 2015). Intine enriched in cellulose, hemicelluloses, pectin and glycoproteins comparable with the primary cell wall of somatic cells. Exine is composed of nexine and sexine, and sexine consists of tectum and baculae (Radja et al., 2019). The pollen coat development largely depends on the tapetum which provides nutrients and exine precursors for the developing pollen. After the microspores are released, tapetal cells constantly synthesize precursors of sporopollenins, the main exine component, and secret them by lipid transfer proteins or ATP-binding cassette transporters (ABCGs) to deposit on the pollen exine. ABCG9 and ABCG31 show high expression in the tapetum, *abcg9* and *abcg31* mutants exhibit immature pollen coat (Choi et al., 2014). ABCG26 and ABCG15 are demonstrated to be important for male reproduction (Quilichini et al., 2014).

Pollen wall development largely depends on efficient secretion of exine precursors in tapetum cells. At late stages of pollen development, tapetal cells synthesize and deposit lipidic materials in the proplastid-derived elaioplasts and the endoplasmic reticulum (ER)-derived tapetosomes (Suzuki et al., 2013). Elaioplasts are rich in steryl esters, free polar lipids and plastid lipid-associated proteins and are filled with globuli (Quilichini et al., 2014). Tapetosomes with high electron dense consist of a fibrous meshwork of vesicles, fibrils and oil bodies with oleosin proteins, alkanes and flavonoids (Hsieh and Huang, 2005). During tapetum degeneration, the contents in elaioplasts and tapetosomes are released and deposited between the exine baculae to complete pollen coat formation (Hsieh and Huang, 2005). Xyloglucan (XyG), the most abundant hemicellulose in primary wall, consists of a *β*-(1,4)-glucan backbone decorated with D-xylosyl chains and is important for the structural organization of the cell wall. The XyG xylosyltransferases (XXTs) catalyze XyG biosynthesis. XXT1 and XXT2 are responsible for initial xylosylation of the glucan backbone (Zabotina et al., 2012). XXT5 likely produces fully xylosylated XyG, XXXG-type XyG by decorating a partially xylosylated glucan backbone produced by XXT1 and/or XXT2 (Zabotina et al., 2008; Culbertson et al., 2018). The cell wall polysaccharides isolated from *xxt5* seedlings contain decreased XyG quantity and reduced glucan backbone substitution with xylosyl residues (Zabotina et al., 2008). XXT5, a Golgi-localized transmembrane protein, catalyzes XyG synthesis by forming protein complexes, XXT2-XXT5 and XXT5-CSLC4. And XXT5 may function as a regulator or an organizer for the XyG synthetic complex (Zabotina et al., 2008; Chou et al., 2012).

Biosynthesis of PT cell wall involves multiple subcellular compartments and vesicle transport pathways. Most of the cell wall polymers such as hemicelluloses and pectin are synthesized at the Golgi apparatus and then secreted at PT tip; and the cell wall glycoproteins such as AGPs and extensins complete their post-translational modification at the Golgi before being exocytosed (Dehors et al., 2019). Other cell wall polymers such as cellulose and callose are synthesized by PM-localized cellulose synthases (CESAs) or callose synthase (CalS)/glucan synthase-like (GSL) complexes (Farrokhi et al., 2006). To date, ten CESA proteins have been identified in *Arabidopsis*. CESA1, CESA3 and CESA6 are involved in cellulose biosynthesis of primary cell wall, while CESA4, CESA7 and CESA8 are active during secondary cell wall establishment (Polko and Kieber, 2019). *cesa1* and *cesa3* mutants are gametophytic lethal. Half of pollen grains from the heterozygous *cesa1+/-* or *cesa3+/-* plants are deformed and ungerminable. While, *cesa6* pollen grains seem normal. However, pollen grains of *cesa2/6/9* triple mutants are deformed and sterile (Persson et al., 2007). CESA6 can be partly complemented by CESA2, CESA5 and CESA9, suggesting partially redundant roles in primary wall biosynthesis (Desprez et al., 2007; Persson et al., 2007). CESA10 function is likely similar to CESA1 based on sequence homology, its biological role remains unclear (Griffiths et al., 2015).

The secretory machinery is responsible for transport of secretory proteins and chemical compounds for maintaining PM homeostasis or cell wall biosynthesis. Coat protein complex II (COPII) vesicles mediate the early secretory pathway, i.e. the newly synthesized proteins and lipids are transported from the ER to the Golgi apparatus where the lipids and proteins receive modification (Hutchings and Zanetti, 2019). The COPII components are essential for plant growth and development. *Arabidopsis* Secretion-associated RAS 1 (SAR1) coordinates with Secretory23A (SEC23A) to control ER export (Zeng et al., 2015). SEC23A and SEC23D localize at the ER exit sites and regulate ER export of proteins and lipids which are necessary for pollen wall formation and exine patterning (Aboulela et al., 2018). SEC24A regulates endomembrane integrity and male fertility (Faso et al., 2009; Conger et al., 2011). SEC24B and SEC24C affect male and female gametophyte development redundantly (Tanaka et al., 2013). SEC31B is required for pollen wall development, probably by regulating the early secretory pathway in tapetal cells. *sec31b* mutant is partially abortive resulted from impaired pollen wall formation (Zhao et al., 2016). SEC31A and SEC31B are redundant in gametogenesis and PT growth (Liu et al., 2021). PT tip growth requires active exocytosis for cell wall biosynthesis and signaling at the apical region (Luo et al., 2017). Thus, PT requires the cargoes to be polarly transported to the apex, such as highly methylesterified pectin (Luo et al., 2017), pectin methyl-esterase (PME) (Wang et al., 2013) etc. The highly methylesterified pectin zone at the tip guides PT elongation and navigates PT to female gametes (Luo et al., 2017; Guo and Yang, 2020). The vesicle trafficking regulators RABA4D (Szumlanski and Nielsen, 2009) and exocyst tethering complex consisting of SEC3, SEC6, SEC8, Exo70A1 (Bloch et al., 2016) and Exo70C2 (Synek et al., 2017) are involved in polar exocytosis.

For the fusion of vesicles with their target membranes, many regulators are involved, such as Rab-GTPases and their effectors, Sec1/Munc18-related (SM) proteins, and soluble *N*-ethylmaleimide-sensitive factor attachment protein receptors (SNAREs) (Ohya et al., 2009; Karnik et al., 2013 Yoon and Munson, 2018). SNARE proteins mediate membrane fusion by forming a trans-SNARE complex, which typically contains four subunits, Qa-, Qb-, Qc-SNAREs on the target membrane, and a R-SNARE on the vesicle (Uemura and Ueda, 2014). Golgi-localized Qc-SNAREs, BET11 and BET12 are required for embryo development and PT development (Bolaños-Villegas et al., 2015). R-SNARE SEC22 regulates vesicle transport between the ER and the Golgi and is involved in pollen development (El-Kasmi et al., 2011). In yeast, syntaxin Sed5p localizes at the cis-Golgi and plays a central role in mediating both anterograde traffic from the ER to the Golgi and retrograde traffic within the Golgi apparatus (Peng and Gallwitz, 2004). Syntaxin5 is the mammalian homologue of Sed5p and potentially regulates the targeting and/or fusion of ER-derived vesicles (Dascher et al., 1994). A plant homolog of Sed5p, Syntaxin of plants 31 (SYP31), localizes at the cis-Golgi and regulates the early secretory pathway (Bubeck et al., 2008). The Golgi resident AtSYP31 is required for anterograde transport from the ER to the Golgi. A recent research demonstrates that AtSYP31 and AtSYP32 are functionally redundant and coordinate with COG3, a subunit of conserved oligomeric Golgi (COG) tethering complex, to regulate intra-Golgi trafficking. The two homolog proteins cooperate to regulate pollen development by controlling protein transport and Golgi morphology (Rui et al., 2020). But the molecular mechanism underlying AtSYP32 regulating pollen development is not completely clear.

In our study, we found that no homozygote progeny was obtained from self-pollinated *atsyp32+/-* alleles due to pollen sterile. Many pollen grains and tubes in *atsyp32+/-*, *AtSYP32* RNAi and *AtSYP3132* RNAi lines burst during germination, and less pollen tubes elongated in pistil and were navigated to the micropyles. The pollen wall structure in these lines were seriously disturbed, and strikingly, there were a large amount of ectopic secretory vesicles retained in pollen cytoplasm. Moreover, the tapetum development and degeneration were severely delayed in the mutants and RNAi lines. Y2H, SLCA and BiFC analysis indicate that AtSYP32 interacted with the PT CWI regulators RALF19 and LRX11, xyloglucan xylosyltransferase XXT5, and the vesicle transport regulators SEC31B, SEC22 and BET12, in the Golgi apparatus. Immunofluorescence assay revealed that the abundance or distribution of PT wall polysaccharides and AGPs altered significantly in the mutants and RNAi lines. Our findings illustrate that AtSYP32 plays essential role in pollen wall development and PT CWI via controlling secretory pathway.

## Results

### *atsyp32* mutants are male sterile

To explore AtSYP32 function on pollen development, we isolated three T-DNA insertion mutants, and confirmed the T-DNA insertion sites by Sanger sequencing (Supplemental Table S1A). The results indicate that in *atsyp32-1*, T-DNA was inserted at the junction site between the second intron and the third exon; in *atsyp32-2*, T-DNA was inserted in the second intron; and in *atsyp32-3*, T-DNA was inserted in the first intron (Figure 1A). Since *AtSYP32* has sequence homology with *AtSYP31*, we also isolated two T-DNA insertion mutants, *atsyp31-1* and *atsyp31-2*, in which the T-DNA cassettes were inserted in the third and the last exons, respectively (Supplemental Figure S1A, Supplemental Table S1B). After screening, we obtained *atsyp31-1* and *atsyp31-2* homozygote alleles, however, we never obtained *atsyp32-1*, *atsyp32-2* and *atsyp32-3* homozygous mutants. We further generated *AtSYP32* RNA interference (*AtSYP32* RNAi), *AtSYP32* overexpressing (*AtSYP32* OE), and *AtSYP31* and *AtSYP32* consensus sequence RNAi (*AtSYP3132* RNAi) lines (Supplemental Table S1, C and D). Meanwhile, we generated anti-AtSYP32 and anti-AtSYP31 polyclonal antibodies to detect their protein abundance. Immunoblot analyses indicate that AtSYP32 endogenous protein levels decreased significantly in *atsyp32-1/-2/-3+/-* mutants, *AtSYP32* RNAi #1/#2 and *AtSYP3132* RNAi #1/#2 lines compared with that in wild type (Figure 1B). In *AtSYP32* OE lines, although endogenous AtSYP32 protein level did not change substantially (Figure 1B), TAP-AtSYP32 (9 *x* myc-AtSYP32) fusion protein was accumulated in a higher level (Figure 1C). The transcription level of *AtSYP32* reduced significantly in *atsyp32+/-* mutants, *AtSYP32* RNAi and *AtSYP3132* RNAi lines compared with that in wild type (Figure 1D, Supplemental Figure S1D). While, the expression of *AtSYP31* in *atsyp31* mutants and *AtSYP3132* RNAi lines reduced dramatically compared with that in wild type (Figure 1D, Supplemental Figure S1B), and the AtSYP31 protein was absent in *atsyp31* mutants (Supplemental Figure S1C). These results indicate that *atsyp32+/-* and *atsyp31* mutants, *AtSYP32* RNAi and *AtSYP3132* RNAi lines were effective for further investigation.

**Figure 1.**
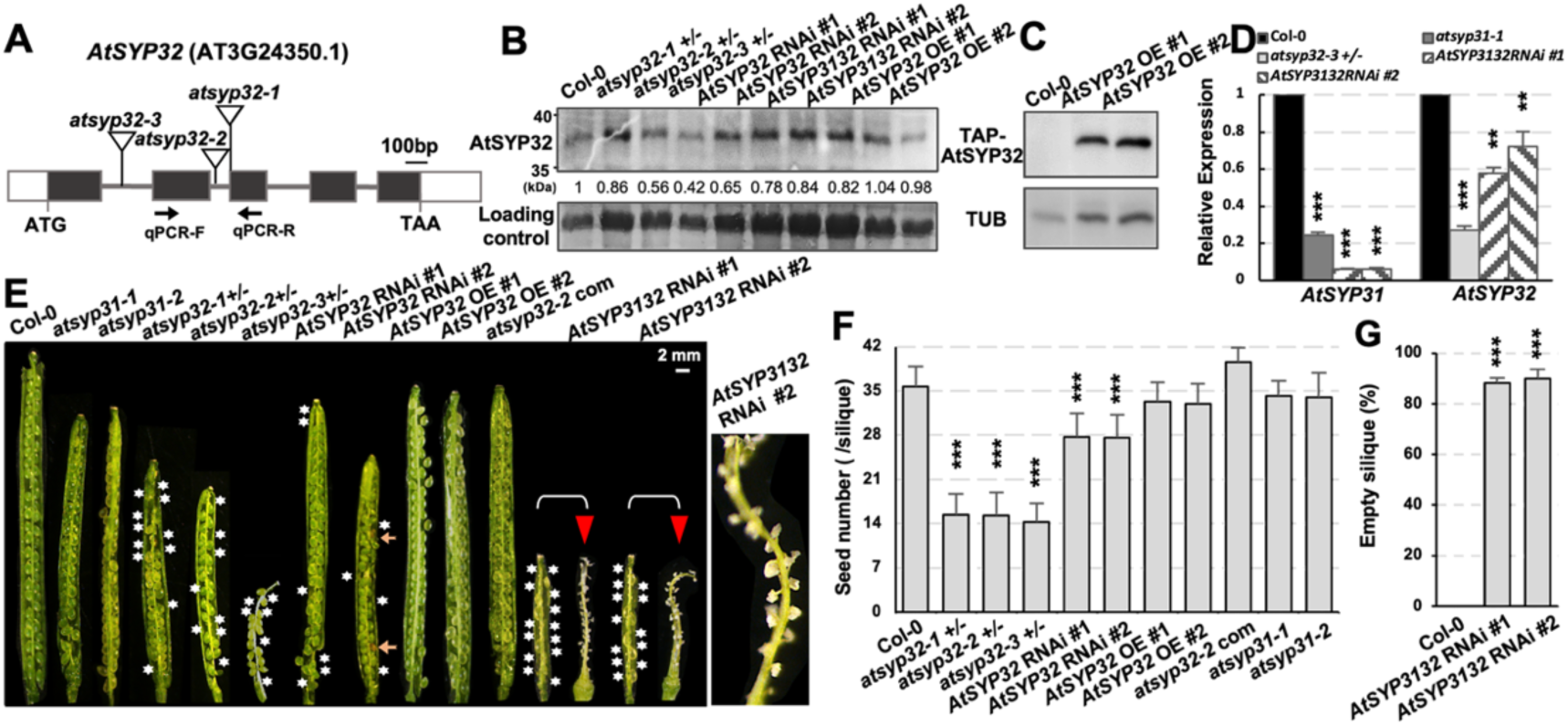
Knockdown of *AtSYP32* led to partial seed sterile. A, Schematic diagrams of *AtSYP32* gene structure and the T-DNA insertion sites (indicated by triangles). Exons are represented by black boxes, introns by solid lines, and untranslated regions by white boxes. Arrows indicate the specific primers for RT-qPCR of *AtSYP32*. B, C, Immunoblot detection of AtSYP32 protein levels with anti-AtSYP32 antibodies (B) and of myc-AtSYP32 protein levels with anti-myc antibody (C) using rosette leaves from 40-day-old plants of the displayed genotypes. The statistics of relative protein levels shown in numbers below the bands were calculated by protein/control band concentrations measured by ImageJ. Coomassie Brilliant Blue (CBB) stained bands of Rubisco served as a loading control in (B); and TUBULIN (TUB) served as an endogenous control in (C). D, Statistics of RT-qPCR determination of relative expression levels of *AtSYP31* and *AtSYP32* in seven-day-old seedlings of the displayed genotypes. Three independent experiments per sample, four technical replicates per experiment. E, Siliques harvested after self-pollination. The magnified panel on right side showing an undeveloped silique with empty seed coats. The white asterisks indicate the sterile seeds; the orange arrows indicate the abnormal seeds; and the red triangles indicate the undeveloped siliques. F, Statistics of seed number per silique of the indicated genotypes. Values are means ± SD (n ≥ 50) from ≥ 3 plants per line. G, Statistics of ratio of undeveloped siliques in wild-type and *AtSYP3132* RNAi lines. Values are means ± SD (n ≥ 100) from ≥ 3 plants per line. **, *P* < 0.01; ***, *P* < 0.001; Student’s *t*-test.

Phenotypic observation indicate that the root length of seven-day-old seedlings of *atsyp32+/-* mutants, *AtSYP32* RNAi and *AtSYP3132* RNAi lines were significantly longer than that of wild type (Col-0), while that of *AtSYP32* OE lines was significantly shorter than that of wild type. The root length of *atsyp31* mutants had no significant difference (Supplemental Figure S1, E and F). The 60-day-old plant height of the *atsyp32+/-*, *AtSYP3132* RNAi and *AtSYP32* OE lines were significantly higher than that of wild type, especially the *AtSYP3132* RNAi lines had the most significance; while, that of *atsyp31* and *AtSYP32* RNAi lines had no obvious difference compared with that of wild type (Supplemental Figure S1G). An important phenotype was that *atsyp32+/-* mutants, *AtSYP32* RNAi and *AtSYP3132* RNAi lines were partially sterile. The siliques of these lines were significantly shorter than those of wild type, and the seed number per silique were significantly less than that in wild type (Figure 1E, 1F). Furthermore, there were many undeveloped siliques in these lines (Supplemental Figure S1H, arrows), especially in *AtSYP3132* RNAi lines, most of the siliques were undeveloped and with empty seed coats (Figure 1, E and G; Supplemental Figure S1H). While, there was no abortion found in *AtSYP32* OE lines and *atsyp31* mutants (Figure 1, E and F). The seed size of *atsyp32+/-* and *atsyp31* mutants, *AtSYP3132* RNAi and *AtSYP32* OE lines seemed larger than those of wild type, especially *AtSYP3132* RNAi lines had large seeds (Supplemental Figure S1I). And the thousand grain weight (TGW) of these lines except for *AtSYP32* RNAi #1, were significantly higher than that of wild type (Supplemental Figure S1J). Moreover, the whole seed protein levels of *atsyp32+/-* mutants, *AtSYP3132* RNAi and *AtSYP32* OE lines were higher than that of wild type, and *atsyp31* mutants and *AtSYP32* RNAi lines had no obviously difference compared with that of wild type (Supplemental Figure S1K). To confirm whether *AtSYP32* is the causal gene for the mutant phenotypes, we generated complementation lines by introducing *pAtSYP32*:*gAtSYP32* (*AtSYP32* genomic fragment driven by 2,000 bp upstream sequence) into *atsyp32-2+/-* mutant (*atsyp32-2* com) via floral dip. In *atsyp32-2* com lines, *AtSYP32* expression was recovered (Supplemental Figure S1D) and the abortive defects was also restored (Figure 1, E and F; 2A-C; Supplemental Figure S1, H and I; S2). All these results indicate that AtSYP32 is essential for plant reproductive development, while AtSYP31 and AtSYP32 are partially functionally redundant, but AtSYP32 plays a predominant role.

**Figure 2.**
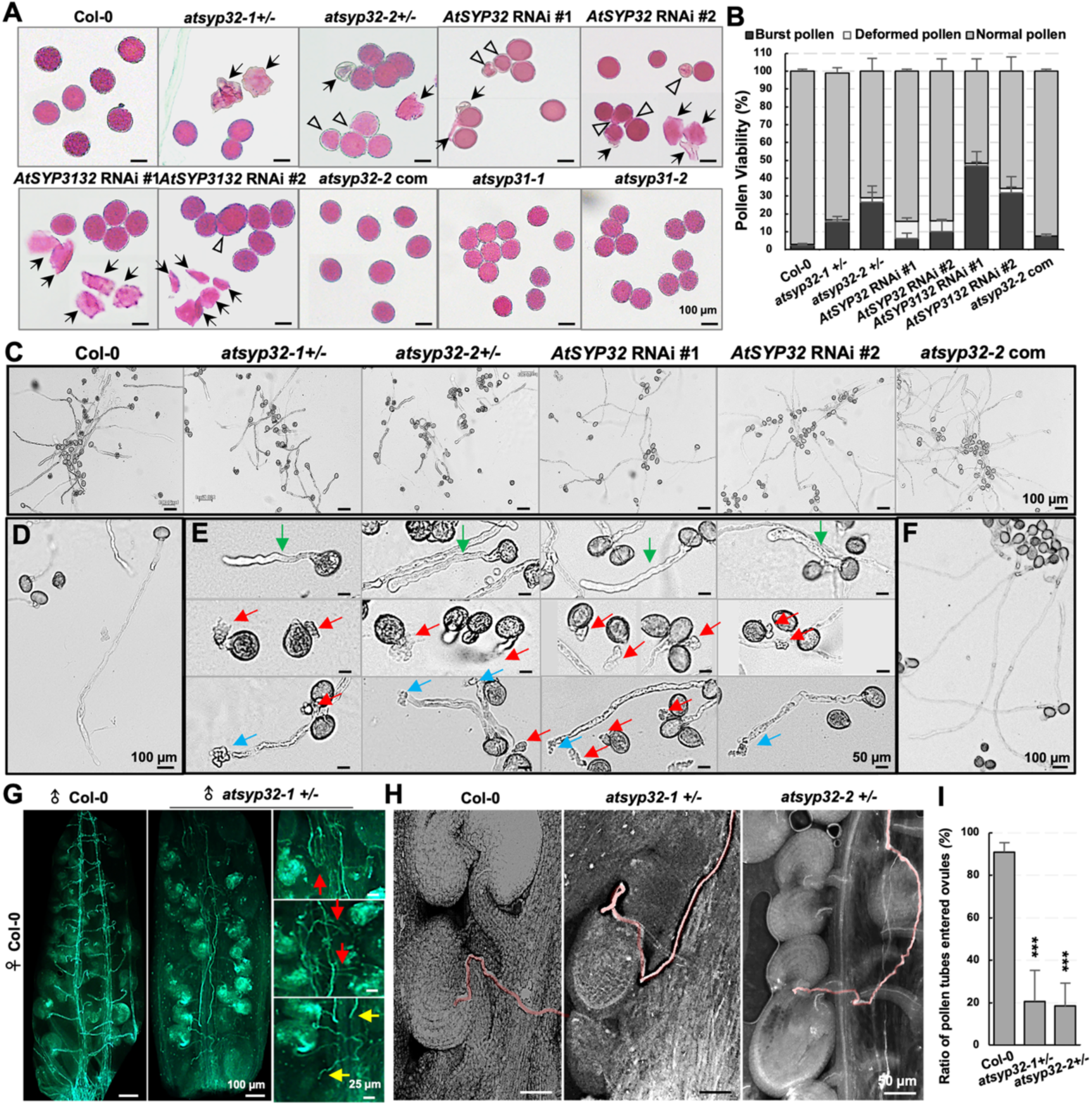
The pollen viability and PT growth in *AtSYP32* knockdown lines. A, Alexander’s staining of pollen grains of the indicated genotypes. The arrows indicate the burst pollen grains, and the triangles indicate the deformed ones. B, Statistics of the pollen viability according to Alexander’s staining results. n ≥ 500. Three biological replicates per sample. C, Pollen grains of Col-0, *atsyp32+/-*, *AtSYP32* RNAi, and *atsyp32-2* com lines germinated *in vitro* for 4h. D-F, The magnified views of pollen grains and tubes. Green arrows, inflated and short PTs; red arrows, burst pollen grains; blue arrows, burst PTs. G, Aniline blue staining of elongated PTs at 48h after pollination on Col-0 pistils with pollen grains of Col-0 and *atsyp32-1+/-*, respectively. Red arrows, not grow straightly PTs; yellow arrows, PTs not enter ovules. H, The PTs entered the ovules (shadowed by red lines). Note that in the mutants, some ovules didn’t have PT entered so kept smaller size. I, Statistics of ratio of PTs entered the ovules. Values are means ± SD (n_mutants_ ≥ 130, n_Col-0_ ≥ 235 ovules in 5 pistils, respectively), three independent experiments per sample. ***, *P* < 0.001; Student’s *t*-test.

### AtSYP32 is essential for pollen viability and pollen tube growth

To clarify the cause of sterile in *atsyp32+/-* mutants, *AtSYP32* RNAi and *AtSYP3132* RNAi lines, we firstly determined whether the transmission of gene mutation was interfered by male gametes or female gametes, via cross-pollination between Col-0 and *atsyp32+/-* mutants. First of all, progeny from self-pollinated *atsyp32-1+/-* and *atsyp32-2+/-* presented the ratio of no mutation : heterozygous : homozygous ≈1:1:0, namely no homozygote was obtained (Table 1). During cross-pollination, when Col-0 served as female parent, the segregation ratio was 107:0 and 82:0, conversely, when *atsyp32-1+/-* and *atsyp32-2+/-* served as female parents, the segregation ratio was 211:69 (roughly 3:1) and 194:42 (roughly 5:1), respectively (Table 1). These results indicate that the *atsyp32* mutations were transmitted by female rather than by male gametophytes. Then, we focused on AtSYP32 regulatory role in the male gametophyte development.

**Table 1.** Segregation of the self-pollinated progenies of *atsyp32+/-* mutants and of cross-pollinated progenies between *atsyp32-1/-2 +/-* and wild type (WT).

| Parental genotype ♀x♂ | Progeny Genotype | Observed Ratio | Approximately Ratio | Expected Ratio |
| --- | --- | --- | --- | --- |
| *atsyp32-1+/-* x *atsyp32-1+/-* | *atsyp32-1+/+ : +/- :-/-* | 152:131:0<sup>a</sup> | 1:1:0 | 1:2:1 |
| *atsyp32-2+/-* x *atsyp32-2+/-* | *atsyp32-2+/+ : +/- :-/-* | 165:153:0<sup>a</sup> | 1:1:0 | 1:2:1 |
| *atsyp32-1+/-* x WT | *atsyp32-1+/+ : +/-* | 211:69<sup>b</sup> | 3:1 | 1:1 |
| *atsyp32-2+/-* x WT | *atsyp32-2+/+ : +/-* | 194:42<sup>b</sup> | 5:1 | 1:1 |
| WT x *atsyp32-1+/-* | *atsyp32-1+/+ : +/-* | 107:0<sup>c</sup> | 1:0 | 1:1 |
| WT x *atsyp32-2+/-* | *atsyp32-2+/+ : +/-* | 82:0<sup>c</sup> | 1:0 | 1:1 |

a, b, c Significance compared with the expected segregation ratio ( $\chi^2$ , $P < 0.01$ ).

To determine pollen viability of the mutants, Alexander’s staining was performed. The results indicated that *atsyp32+/-* mutants, *AtSYP32* RNAi and *AtSYP3132* RNAi lines had less viable microspores in the anthers compared with wild type; especially in *AtSYP3132* RNAi #1/#2 lines, only a few pollen grains were viable (Supplemental Figure S2A). Moreover, many pollen grains in these lines were ruptured (Figure 2A, arrows). The burst ratio of pollen grains in *atsyp32-1/-2+/-* mutants were about 16% and 27%, and in *AtSYP3132* RNAi #1/#2 lines reached about 47% and 32%, respectively, compared with 0.64% in wild type (Figure 2B). Moreover, there are many deformed pollen grains (Figure 2A, triangles). In particular, the deformity ratio in *AtSYP32* RNAi #1/#2 lines reached about 10% and 6%, respectively (Figure 2B). DAPI staining indicated that in contrast to wild-type pollen grains with two or three nuclei, part of *atsyp32+/-* and *AtSYP32* RNAi pollen grains had no DAPI signals (Supplemental Figure S2B, arrows and asterisks), the ratio of microspores with nuclei were significantly lower than that in wild type (Supplemental Figure S2C), suggesting that meiosis was affected in these lines. These phenotypes indicate that AtSYP32 is essential for pollen development and viability.

To test whether the pollen tube (PT) performance in these mutants was affected, an *in vitro* pollen germination assay was performed. After 4h incubation, the pollen germination ratio of *atsyp32-1/-2+/-* were 62% and 56%, and that of *AtSYP32* RNAi #1/#2 lines were 61% and 64%, respectively, which were significantly lower than 76% of wild type (n≈500, three repeats) (Supplemental Figure S2D). Moreover, many shorter, wavy and inflated PTs were observed, especially in *atsyp32-1/-2+/-* mutants, most of PTs were significantly shorter than those in wild type and *AtSYP32* RNAi lines (Figure 2, C-E). Moreover, many pollen grains and tubes were ruptured in *atsyp32+/-* mutants and *AtSYP32* RNAi lines (Figure 2E, arrows). While, these phenotypes were restored in *atsyp32-2* com lines (Figure 2, C and F), indicating that *AtSYP32* is critical for integrity of pollen grains and PTs.

The PT growth were further validated *in vivo*. We pollinated wild-type stigma with Col-0 and *atsyp32-1+/-* pollen grains and examined the elongation. At 48 h after pollination, the wild type PTs labeled by Aniline blue dyeing elongated straightly, and almost every ovule had a PT realize micropyles and completed double fertilization that visualized by expanded ovules. While, elongated *atsyp32-1+/-* PTs were much less than those of wild type, and many PTs did not elongate straightly or enter ovules (Figure 2G, arrows), and many ovules didn’t have PTs entered to complete double fertilization (Figure 2, G and H), so the ratio were significantly reduced compared with that of wild type (Figure 2I). Collectively, the *in vitro* and *in vivo* assay indicate that *AtSYP32* is crucial for pollen viability, PT growth and targeting to the ovule.

### AtSYP32 is required for pollen wall development via controlling secretion pathway

Since pollen hydration on the stigma is a critical step for pollen germination and PT growth (Li et al., 2017), we checked pollen hydration, and found that wild-type and part of *atsyp32-1+/-* pollen grains got hydrated on Col-0 stigma within 5 min (Figure 3, A and B, arrows), while part of *atsyp32-1+/-* pollen grains did not hydrate even after 10 min (Figure 3B, arrow heads). Since pollen wall is essential for hydration (Zhan et al., 2018), scanning electron microscope (SEM) analysis was performed to observe pollen wall structure. Compared with wild-type plump pollen grains with typical reticular pollen wall, some pollen grains of *atsyp32-1+/-* and *atsyp32-2+/-* were shriveled or exhibited deformed morphology (Figure 3, C and D, arrow heads); even more, the pollen wall structures were incomplete (Figure 3E, arrows). The deformity ratio (including incomplete pollen wall) in *atsyp32-1+/-* and *atsyp32-2+/-* were 17% and 8%, respectively, which were significantly higher than % of wild type (Figure 3F). Transmission electron microscope (TEM) analysis was performed to observe the ultrastructure of pollen wall. Compared with wild-type intact and even pollen wall structure, many pollen grains in *atsyp32+/-*, *AtSYP32* RNAi and *AtSYP3132* RNAi lines was disordered and incomplete (Figure 3G). Moreover, many pollen grains were ruptured or deformed, and some of them were adhere to the tapetum or to each other (Figure 3G, arrows; Supplemental Figure S3A). The intine in *atsyp32+/-* mutants and *AtSYP32* RNAi lines were significantly thinner than that of wild type (Figure 3H, in between two red triangles; 3K); and thickened and formed a striped structure at some places (Figure 3Ik; Supplemental Figure S3Cm, cruciform stars). On the other hand, the sexine was irregularly organized and unevenly distributed on the nexine (Figure 3H, arrows; Supplemental Figure S3B). And in *AtSYP3132* RNAi lines, it was hard to distinguish the intine, and the organization of exine was also disrupted (Figure 3Hf, Supplemental Figure S3Be). The ratio of microspores with defective pollen wall was significantly higher than that of Col-0 (Supplemental Figure S3D, deep gray columns). These phenotypes suggest that AtSYP32 regulates pollen wall formation. A remarkable phenotype was that in *AtSYP3132* RNAi lines, about 31% microspores had no cytoplasmic structure (Supplemental Figure S3D, ‘empty pollen grains’), about 59.5% exhibited plasmolysis (‘plasmolytic pollen grains’), and only 9.5% had cellular structures (‘cytoplasmic pollen grains’). And in *atsyp32-2+/-* mutant, there was 2.6% microspores had no cytoplasmic structure (Supplemental Figure S3D). These phenotypes indicate that AtSYP32 is required for pollen and pollen wall development; AtSYP31 and AtSYP32 were functionally redundant, and AtSYP32 played predominate roles.

**Figure 3.**
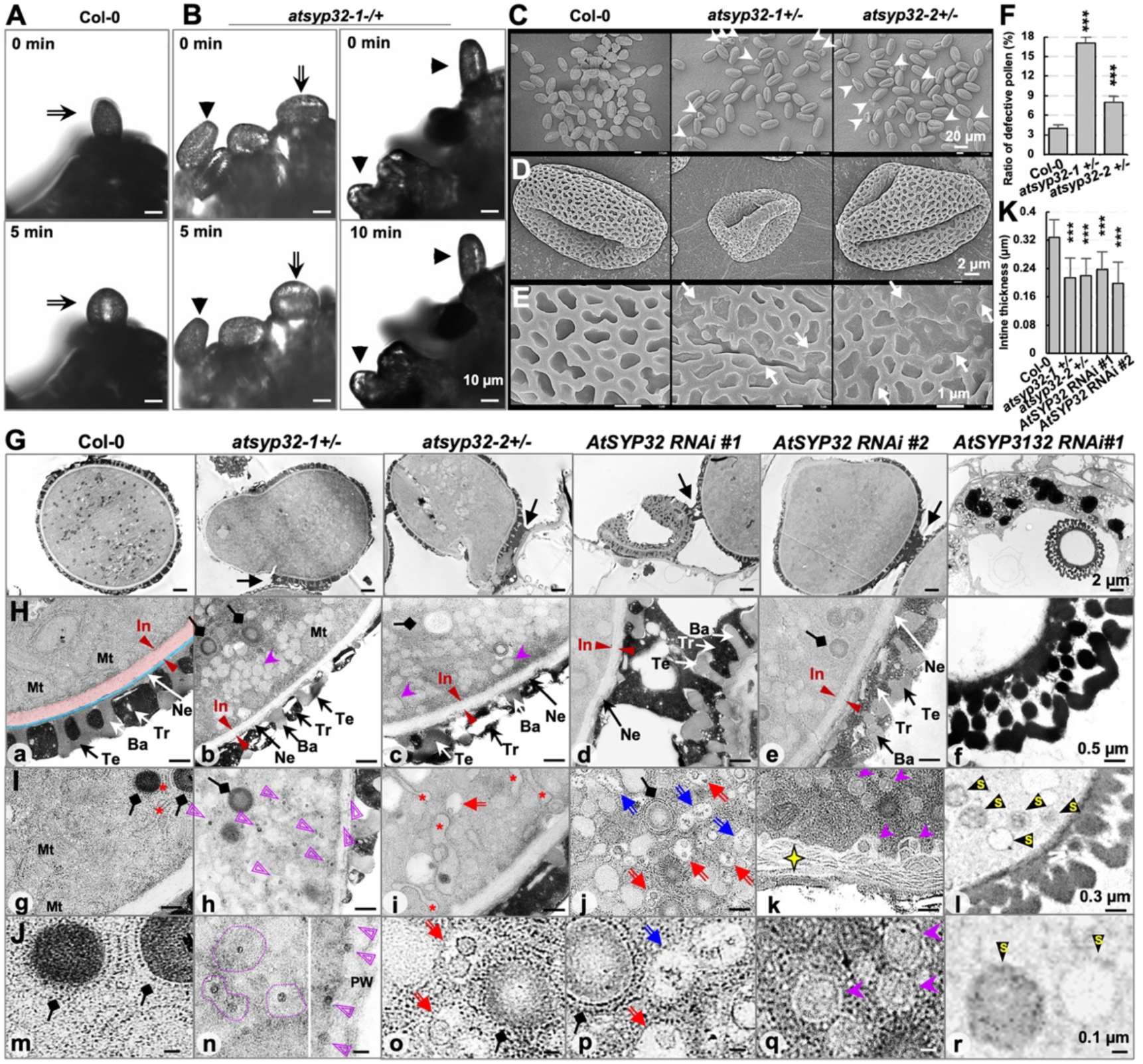
The pollen defects in *AtSYP32* knockdown lines. A, B, Pollen hydration was detective in *atsyp32-1+/-*. Pollen grains from *atsyp32-1+/-* and Col-0 were pollinated on Col-0 stigmas, respectively. The arrows indicate the hydrated pollen grains from Col-0 and *atsyp32-1+/-*. The triangles indicate the un-hydrated *atsyp32-1+/-* pollen grains. C-E, Scanning electron microscopy (SEM) observation of pollen wall of Col-0, *atsyp32-1+/-* and *atsyp32-2+/-*. Arrowheads indicate the deformed pollen grains, and arrows indicate the fractured pollen wall. F, Statistics of ratio of defective pollen shown representatively in (C). Values are means ± SD (n ≥ 500), three biological replicates per sample. G, Ultrastructure of pollen grains. Arrows label the adhesion between pollen grains or pollen grains and epidermal cells. H, Ultrastructure of pollen wall. The structure between the two red arrows is the intine, which is also highlighted by a red shadow in (a), and a blue shadow highlights the nexine. Ba, baculum; In, intine; Mt, mitochondria; Ne, nexine; PW, pollen wall; Te, tectum; Tr. tryphine. I, J, Ultrastructure of the cytoplasm. Diamond arrows, unknown spheroid structures with rough ER; asterisks, rough ER; cruciform stars, thickened intine; red arrows, MVBs; blue arrows, contractile vacuoles; purple dovetail arrow heads, EXPOs; black triangles with yellow letter ‘s’, SVs; purple triangles and circles, unknown structures with tiny and high electron dense core. Mt, mitochondria. K. Statistics of intine thickness of the displayed lines. Values are means ± SD (n ≥ 10) from 3 biological replications. ***, *P* < 0.001; Student’s *t*-test.

In wild-type pollen grains, the ER mainly presented a tubular structure, while in the *atsyp32-2+/-* and *AtSYP32* RNAi lines, expanded tubular or vesicular structures with ribosomes, namely inflated ER, were observed (Figure 3I; Supplemental Figure S3B and C, red asterisks), suggesting that AtSYP32 is required for ER morphology maintenance. Another striking feature was the numerous vesicles appeared in pollen cytoplasm in *atsyp32+/-* mutants and *AtSYP32* RNAi lines rather than in wild type (Figure 3, H-I). Firstly, some of the vesicles may be multivesicular bodies (MVBs) (Figure 3, H-J; Supplemental Figure S3C, red arrows) and contractile vacuoles (blue arrows), some likely the exocyst positive organelles (EXPOs) (purple dovetail arrow heads) which mediate the release of a cytosol-containing exosome to the apoplast (Wang et al., 2010), and some look like secretory vesicles (SVs) (black triangles with letter ‘s’) which deliver cargoes from the TGN to the PM/apoplast (van de Meene et al., 2017). On the other hand, there were some unknown structures. A structure with tiny and high electron dense core (Figure 3, Ih and Jn, purple triangles and circles) retained in the mutants and RNAi lines rather than in wild type. Some of them were next to the plasma membrane (PM), and some of them were in the pollen wall (Figure 3, Ih and Jn, purple triangles), likely the cores were being secreted to the pollen wall. Moreover, some other unknown structures with high electron dense (Supplemental Figure S3B, yellow arrows), or with either large or small dense cores (Supplemental Figure S3C, blue triangles) were also observed. In addition, a spherical structure with high electron density and surrounded by rough ER was observed in wild-type pollen cytoplasm (Figure 3, G-J, Supplemental Figure S3, B and C, black diamond arrows); while, in *atsyp32+/-* mutants and *AtSYP32* RNAi lines, number of this structure reduced substantially (Figure 3G), and the electron density was lower than that in wild type (Figure 3J). Collectively, the inflated ER, MVBs, EXPOs, SVs and the unknown structures found in *atsyp32+/-* mutants, *AtSYP32* RNAi and *AtSYP3132* RNAi lines indicate that vesicle trafficking was blocked seriously. These results suggest that AtSYP32 controls secretion pathway which is responsible for pollen wall biosynthesis.

### AtSYP32 is critical for tapetum development and degeneration

The phenotypes of ruptured pollen grains and tubes, incomplete pollen wall and delayed PT growth strongly indicated an abnormal deposition of pollen wall in *atsyp32+/-* mutants, *AtSYP32* RNAi and *AtSYP3132* RNAi lines. Aniline blue staining assay indicated an ectopic callose deposition in pollen grains in these lines (Supplemental Figure S4, A and B), and the ratio were significantly higher than that in wild type (Supplemental Figure S4C), validating the abnormality of pollen wall deposition.

Since the pollen coat formation is mainly dependent on tapetum (Lou et al., 2018), we observed the tapetum structure. In wild type, the tapetum at the late uninucleate stage (Figure 4Aa, pink shadow) is characterized by the formation of two specialized storage organelles, tapetosomes with high electron dense (Figure 4B, arrows) and the elaioplasts with electron-lucent plastoglobules (Figure 4B, arrow heads). While, the tapetum in *atsyp32-1+/-*, *AtSYP32* RNAi #1 and *AtSYP3132* RNAi #1 lines at the same stage were much thinner than that in Col-0 (Figure 4A), and some tapetal cells had enlarged vacuole (Figure 4Ab, asterisk). At the bicellular stage, the wild-type tapetal cells degenerated and bicellular microspores developed (Figure 4Ce). By contrast, in *atsyp32+/-, AtSYP32* RNAi, and *AtSYP3132* RNAi lines, the tapetum degenerated unevenly and disconnected at some places. Some tapetosomes seemed to fuse with each other to form a large tapetosome; and the amount of elaioplasts likely decreased compared with that in Col-0 (Figure 4C). At the tricellular stage, wild-type tapetum completely degenerated (Figure 4Di). However, in the mutants and RNAi locules, lots of tapetum residues remained (Figure 4, Dj-l, orange arrows), indicating a significant delay in tapetum degeneration. These findings suggest that AtSYP32 regulates tapetum development and degeneration which is critical for pollen wall development.

**Figure 4.**
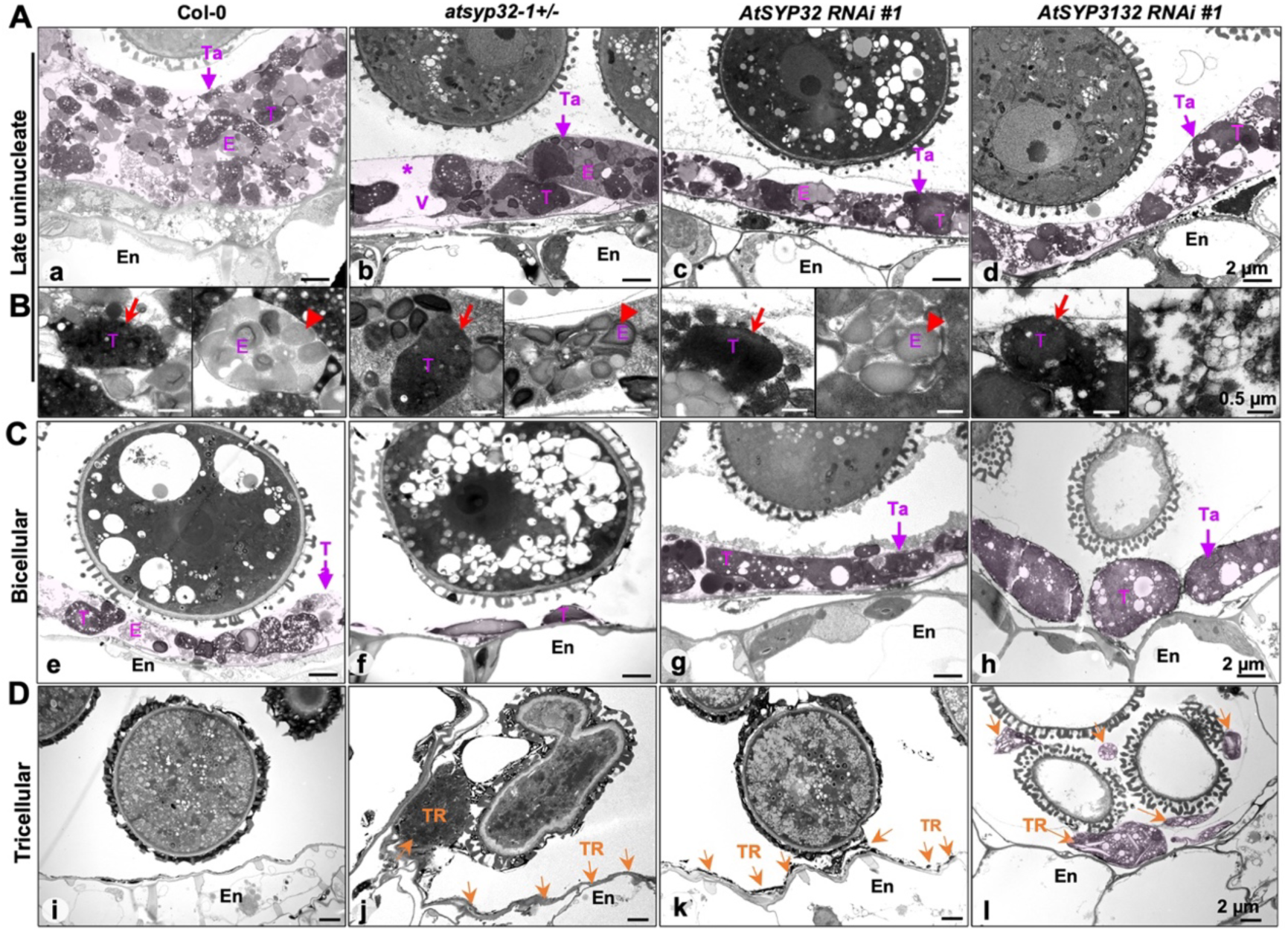
Ultrastructure of tapetum at different developmental stages. A, B, Ultrastructure of tapetum at late uninucleate stage. Magnified images of tapetosomes (red arrows) and elaioplasts (red triangles) are shown in (B). Asterisk, enlarged vacuole. C, D, Ultrastructure of tapetum at bicellular stage (C) and tricellular stage (D). The tapetum was labeled by pink shadow. Red arrows, tapetosomes; red triangles, elaioplasts; pink arrows, tapetum; orange arrows, tapetum residues. E, elaioplast; En, endodermis; T, tapetosome; Ta, tapetum; TR, tapetum residue; V, enlarged vacuole.

Since AtSYP32 may regulate the development of tapetum, we determined AtSYP32 tissue distribution. Confocal images revealed that in the anthers of mCherry-AtSYP32-expressing line, mCherry-AtSYP32 showed strong signals in the tapetum (Figure 5A). Further observation found that mCherry-AtSYP32 was localized in the pollen grain (Figure 5B) and the PT tip (Figure 5C). This emphasized the close relevance of AtSYP32 with tapetum and pollen wall development, and PT growth.

**Figure 5.**
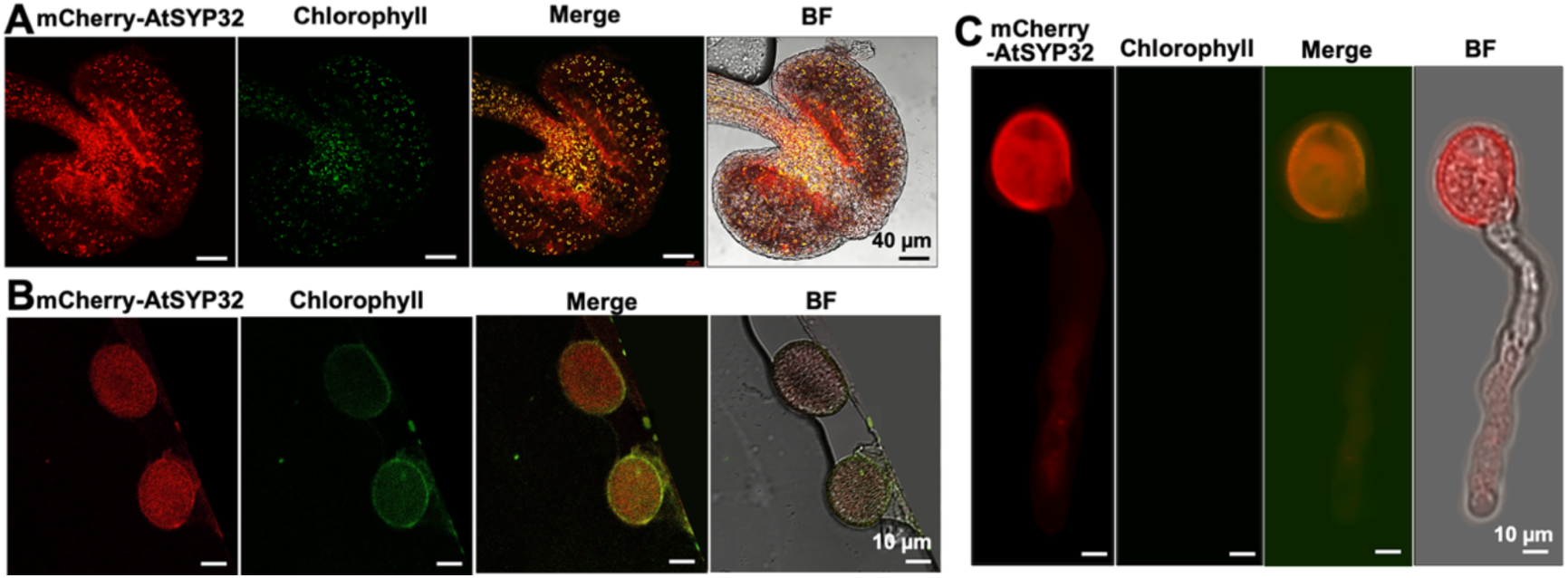
AtSYP32 distribution in tapetum and pollen. mcherry-AtSYP32 is localized at tapetum (A), pollen grain (B) and the PT tip (C). Chlorophyll represents the autofluorescence. BF, bright field.

### AtSYP32 is required for PT cell wall deposition

To investigate the effects of knockdown of *AtSYP32* on PT cell wall, we examined the contents of cell wall polysaccharides and glycoproteins. An immunofluorescence assay was employed to detect the contents of highly methylesterified homogalacturonans (HGs) using anti-JIM7, de-esterified HGs using anti-JIM5, XXXG XyG using anti-LM15, and a subset of arabinogalactan proteins (AGPs) using anti-LM2 antibodies, respectively. In Col-0, the highly methylesterified HGs were distributed in the PT apex; while in *atsyp32-1+/-*, it lost the apex-localization and irregularly distributed, and some of the HGs aggregated inside the PTs (Figure 6A), leading to an significant increase of the contents compared with wild type (Figure 6F). The de-esterified HGs and XXXG XyG were uniformly distributed in PT cell wall in wild type, while in *atsyp32-1+/-*, they distributed disorderly, and some of them were retained inside the PTs, but the content didn’t alter significantly (Figure 6, B, C and F). However, the contents of AGPs labeled by anti-LM2 antibody reduced dramatically in *atsyp32-1+/-* compared with that in wild type (Figure 6, D and F). The pectins are the major components of PT cell wall. In wild-type, Ruthenium red labeled the methylesterified pectins concentrated at PT tips in addition to cell wall distribution, whereas in *atsyp32-1+/-*, the pectins lost polarity distribution at the tips, and can be detected inside the ruptured PTs (Figure 6E), suggest that the pectins were not secreted yet. We further determinated the total cellulose contents of stems. As expected, stem cellulose abundance in *atsyp32+/-*, *AtSYP32* RNAi and *AtSYP3132* RNAi lines decreased significantly compared with that in wild type (Figure 6G), indicating the cellulose biosynthesis was also disrupted. These results imply that AtSYP32 is crucial for secretion and efficient polar transport and polarity maintenance of PT cell wall components.

**Figure 6.**
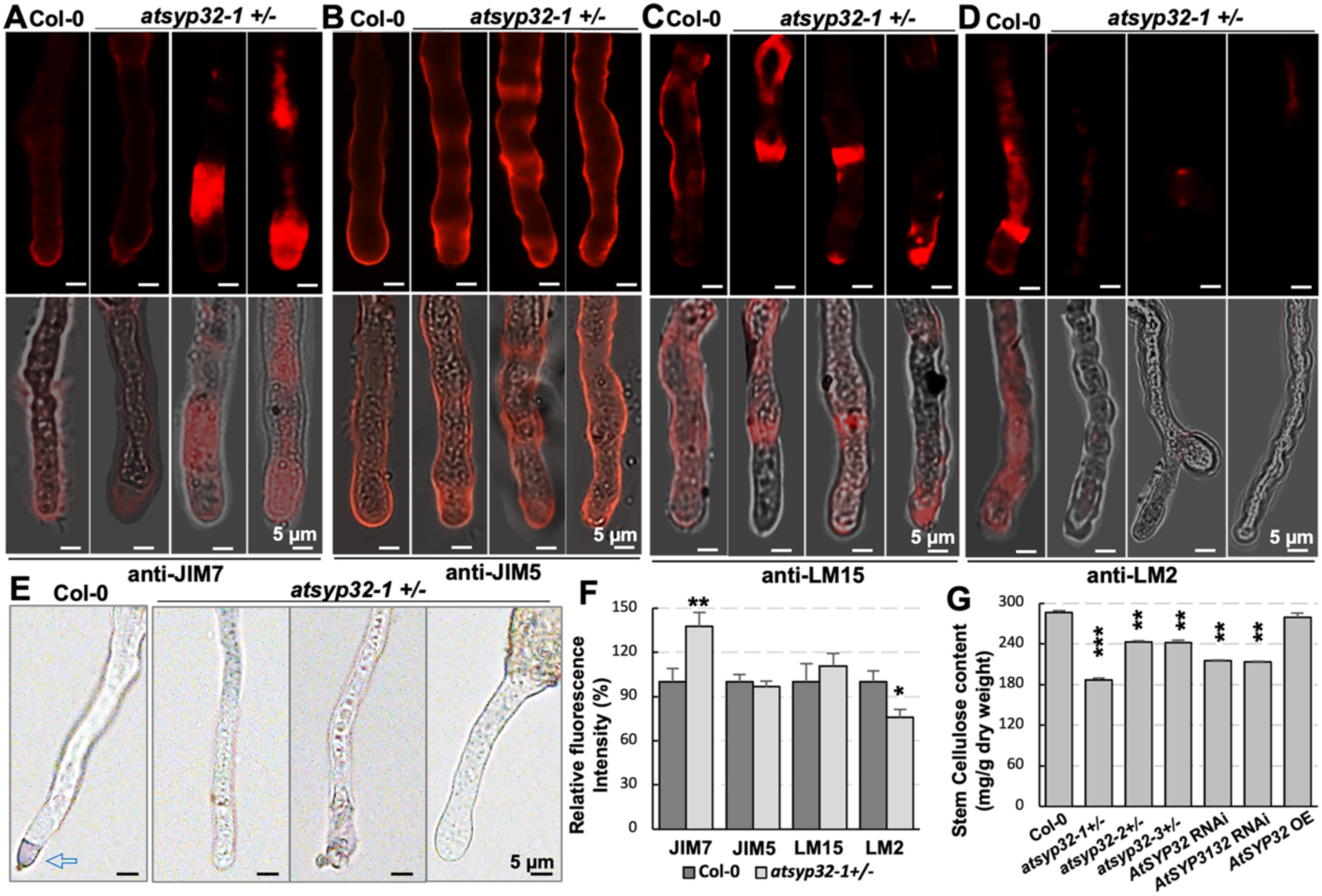
Ectopic deposition of components of PT cell wall in *atsyp32-1+/-* mutant. A-D, Immunolabeling of highly methylesterified HGs with anti-JIM7 antibody (A), de-esterified HGs with anti-JIM5 antibody (B), XXXG xyloglucan with anti-LM15 antibody (C), and AGPs with anti-LM2 antibody (D) in PTs of Col-0 and *atsyp32-1+/-*. The lower panels are the merge of bright felid with fluorescence. E, Pectin detection by Ruthenium red staining of Col-0 and *atsyp32-1+/-* PTs. F, Statistics of relative fluorescence intensities of PTs shown representatively in A-D. Values are means ± SD (n ≥ 10), three biological replicates per sample. G. The stem cellulose contents of wild-type, *atsyp32+/-*, *AtSYP32* RNAi, *AtSYP3132* RNAi and *atsyp32-2* OE lines. Values are means ± SD, three independent experiments per sample. **, *P* < 0.01; ***, *P* < 0.001; Student’s *t*-test.

### AtSYP32 may regulate cell wall biosynthesis and PT CWI via controlling vesicle trafficking

To clarify AtSYP32 regulatory role in PT CWI, we identified AtSYP32-relating factors. Firstly, we searched *AtSYP32*-coexpression genes in ATTED II database (https://atted.jp/) and found 12 pollen tube growth- and cell wall-related genes (Table 2). RT-qPCR determination using total RNA from flowers (pistil removed) indicate that most of them altered significantly in *atsyp32+/- and AtSYP32* RNAi lines compared with that in wild type (Figure 7A). Among them, three genes are XXT5 catalyzes xylosylation of XyG, the most abundant hemicellulose of primary cell walls (Culbertson et al., 2018); Golgi-localized glucuronokinase G (GlcAK) regulates synthesis of UDP-GlcA, the immediate precursor of monosaccharides (D-galacturonic acid, D-xylose and D-apiose) required for cell wall polysaccharide biosynthesis (Borg et al., 2021); and UDP-glucose dehydrogenase2 (UGD2) is involved in biosynthesis of nucleotide sugars as precursors of primary cell wall (Reboul et al., 2011). The expression levels of *XXT5* and *UGD2* reduced significantly in *atsyp32+/-* mutants and *AtSYP32* RNAi lines, while *GlcAK* increased significantly only in *atsyp32-1+/-* compared with that of wild type (Figure 7A), suggesting the cell wall biosynthesis was affected. SEC22 is demonstrated to be crucial for male gametophyte development (El-Kasmi et al., 2011) and *SEC22* expression increased significantly in the mutants and RNAi lines compared with that in wild type (Figure 7A), validating that SEC22 is involved in pollen development. It is well known that RAB proteins play essential roles in membrane trafficking from the trans-Golgi network (TGN) to the PM/cell wall or to the cell plate during cytokinesis, such as RABA4A (Lycett, 2008; Lunn et al., 2013), RAB1A (Peng et al., 2011) and RABA4B (Preuss et al., 2004) (Table 2). VPS45 (Tanaka et al., 2013; Matsuura et al., 2020) and VPS46 (Spitzer et al., 2015) are implicated in post-Golgi trafficking; DL1C is essential for PM maintenance during pollen maturation (Kang et al., 2003); and VAMP721 is involved in secretory pathway and contribute to cell plate formation during cytokinesis (Zhang et al., 2021) (Table 2). Relative expression of these genes altered significantly in *atsyp32+/-* mutants and *AtSYP32* RNAi lines compared with those in wild type (Figure 7A), suggesting AtSYP32 participates the PT cell wall formation-related vesicle trafficking.

**Figure 7.**
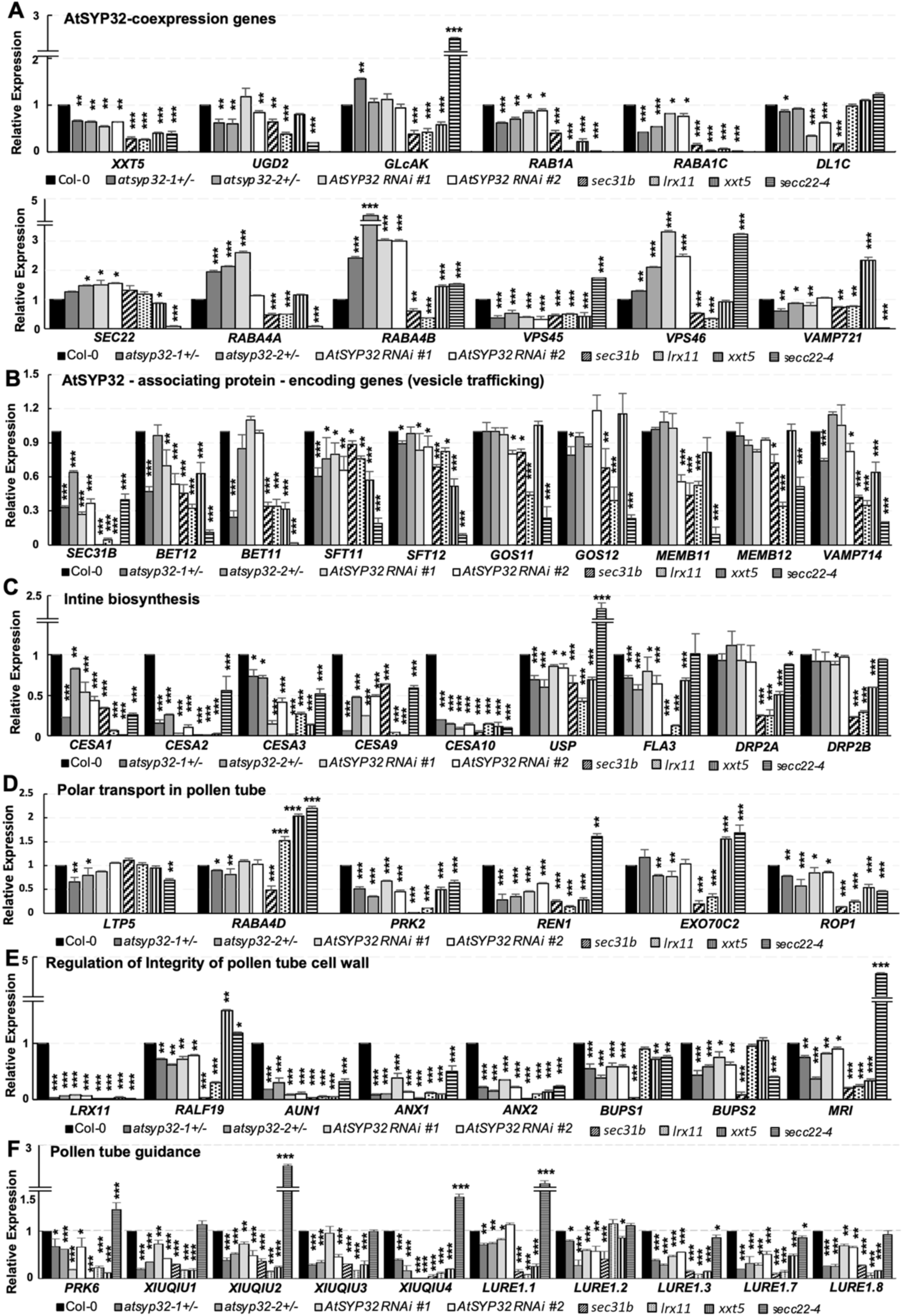
Determination of expression levels of the genes related to pollen wall biosynthesis and PT CWI maintenance. The relative expression of the *AtSYP32*-coexpressing genes (A), the associating protein-encoding genes (B), intine/primary wall biosynthesis-relating genes (C), the polar transport-regulating genes (D), PT CWI maintenance-relating genes (E and F) were determined by RT-qPCR. *ACT2* was used as an endogenous control. In A-E, Total RNA was extracted from the flowers without stigma. In F, total RNA was extracted from the whole flowers. Three independent experiments per sample, four technical replicates per experiment. *, *P* < 0.05; **, *P* < 0.01; ***, *P* < 0.001; Student’s *t*-test.

**Table 2.** *AtSYP32* co-expressional genes.

| Gene ID | Gene name | Function | Support ability | ath-u.2 / SYP32 | ath-r.5/ SYP32 | ath-m.9 / SYP32 | ath-m.4.tis /SYP32 | ath-m.4.str / SYP32 | ath-m.4.hor/ SYP32 | ath-m.4.bio / SYP32 | ath-m.4.lig/ SYP32 | References |
| --- | --- | --- | --- | --- | --- | --- | --- | --- | --- | --- | --- | --- |
| AT1G74380 | *XXT5* | **Catalyzes** xylosylation of XyG, the most abundant hemicellulose of primary cell walls. | 3 | 5.5 | 5.7 | 3.8 | 2.7 | 4 | 3.6 | 4.6 | 1.5 | Chou et al., 2015; Culbertson et al., 2018. |
| AT3G01640 | *GLCAK* | **Regulates** synthesis of UDP-GlcA, the immediate precursor of monosaccharides required for cell wall biosynthesis | 2 | 4.3 | 5.4 | 1.9 | 1.9 | 2.8 | 3.1 | 3.3 | -0.6 | Muthana et al., 2015. |
| AT3G29360 | *UGD2* | **Involved** in biosynthesis of nucleotide sugars as precursors of primary cell wall. | 3 | 5.6 | 6 | 3.6 | 4 | 2.2 | 1.8 | 3.3 | -0.3 | Reboul et al., 2011. |
| AT5G45750 | *RABA1C* | **Implicated** in delivery of cell wall synthetic materials and secretory proteins to the apex in growing root hairs and PTs. | 1 | 5.5 | 5.4 | 4 | 2.3 | 3.6 | 3.8 | 3.1 | 1.6 | Lycett, 2008; Qi and Zheng, 2013. |
| AT5G47200 | *RAB1A* | **Facilitates** membrane trafficking for PT growth. | 3 | 5.9 | 5.8 | 4.4 | -0.3 | 1.4 | 1.4 | 1.1 | 0.7 | Lycett, 2008; Peng et al., 2011. |
| AT5G65270 | *RABA4A* | **Regulates** vesicle trafficking to the cell wall. | 2 | 6 | 5.2 | 5 | 0.4 | 2.7 | 2.3 | 2.5 | 1.2 | Lycett, 2008. |
| AT4G39990 | *RABA4B* | **Implicated** in polarized secretion of cell wall components in plant cells. | 2 | 6 | 5.9 | 4.4 | 1.3 | 2.3 | 2.6 | 3.8 | 1.4 | Preuss et al., 2004; Lycett, 2008. |
| AT1G04750 | *VAMP721* | **Involved** in secretory pathway and contribute to cell plate formation during cytokinesis. | 3 | 5 | 5.1 | 3.4 | 0 | 0.1 | 0.1 | -0.2 | -0.2 | Zhang et al., 2021. |
| AT1G77140 | *VPS45* | **Implicated** in early endocytic route which is crucial for cell polarity and plant architecture. | 3 | 7.6 | 7.2 | 5.9 | 0.6 | 2.4 | 2.8 | 4.7 | 0.6 | Tanaka et al., 2013; Matsuura et al., 2020. |
| AT1G17730 | *VPS46* | **Mediate** MVB sorting of auxin carriers and essential for embryo development. | 2 | 5.5 | 5.3 | 4 | 1.3 | 0.7 | 0.8 | 3.2 | 0.2 | Spitzer et al., 2015. |
| AT1G14830 | *DL1C* | **Essential** for PM maintenance during pollen maturation. | 3 | 6.6 | 6.3 | 5.1 | 2.2 | 1.8 | 2.9 | 2.6 | 0 | Kang et al., 2003. |
| AT1G11890 | *SEC22* | **Crucial** for pollen development and cytoskeleton dynamics. | 2 | 7 | 6.7 | 5.3 | 4.4 | 2.7 | 5.7 | 1.9 | 0.8 | El-Kasmi et al., 2011; Guan et al., 2021. |

Abbreviations: ath-u, microarray-based and RNAseq-based coexpression; ath-r, RNAseq-based coexpression; ath-m, microarray-based coexpression; tis, tissue experiment; str, abiotic stress experiment; hor, hormone experiment; bio, biotic stress; lig, light experiment; DL1C, dynamin-like 1C; GLcAK, glucuronokinase G; RAB1C, Ras-related brain GTPases 1C; SEC22, secretion 22; UGD2, UDP-glucose dehydrogenase 2; VAMP721, vesicle-associated membrane protein 721; VPS45, vacuolar protein sorting 45; XXT5, xyloglucan xylosyltransferase 5.

Then, we identified 17 AtSYP32-associated proteins via pull down assay followed by LC-MS/MS using *myc*-*AtSYP32*-expressing plants, and ten of them were vesicle trafficking regulators (Table 3) among which three factors, SEC22, SEC31B and BET11, are related to pollen wall development. It is worth noting that *SEC22* which is co-expressed with *AtSYP32* was also in this list, i.e. SEC22 is also associated with AtSYP32. In addition, as mentioned above, the COPII coatomer SEC31B regulates pollen wall development by modulating secretory pathway in tapetum (Zhao et al., 2016; Liu X. et al., 2021); and the Qc-SNAREs BET11 and BET12 are required for fertility and pollen tube elongation (Bolaños-Villegas et al., 2015). The expression of *SEC31B* and *BET12* reduced significantly in *atsyp32+/-* mutants and *AtSYP32* RNAi lines, while that of *BET11* reduced significantly only in *atsyp32-1+/-* compared with that in wild type (Figure 7B). These results suggest a potential functional relevance between AtSYP32 and SEC22, SEC31B or BET11/12, respectively. Moreover, the expression levels of *SFT11*/*12* and *VAMP714* decreased significantly compared with that in wild type (Figure 7B), suggesting that AtSYP32-mediated vesicle trafficking was deeply involved in the pollen wall development and PT CWI maintenance.

**Table 3.**
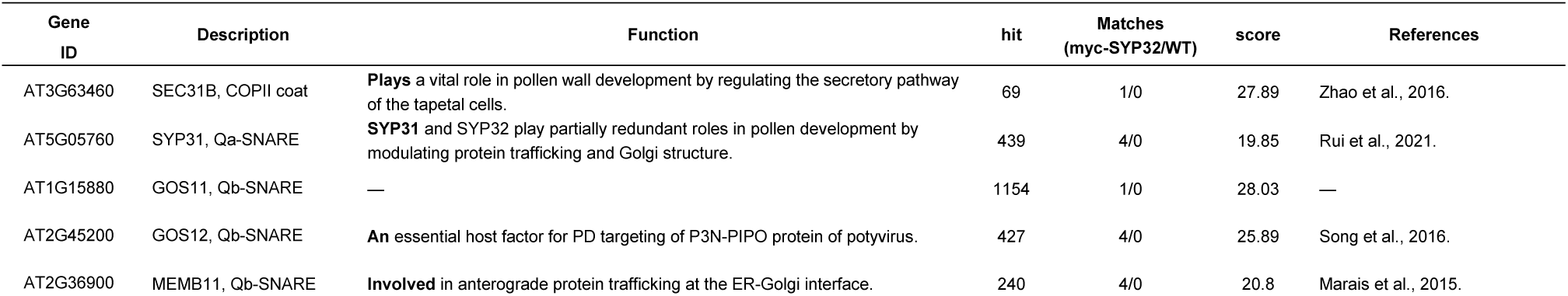
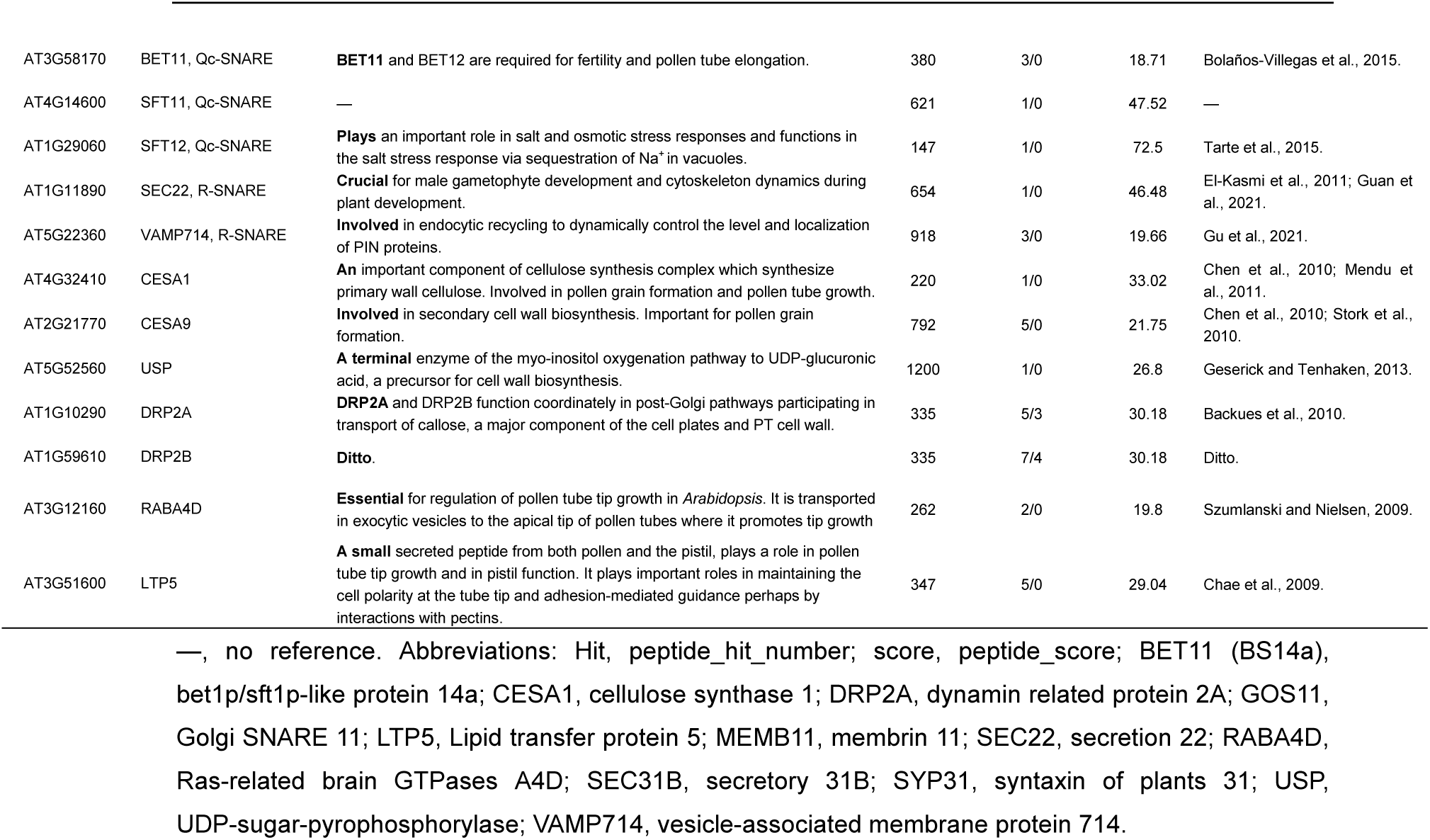
LC-MS/MS identified AtSYP32-associating proteins.

| Gene ID | Description | Function | hit | Matches (myc-SYP32/WT) | score | References |
| --- | --- | --- | --- | --- | --- | --- |
| AT3G63460 | SEC31B, COPII coat | **Plays** a vital role in pollen wall development by regulating the secretory pathway of the tapetal cells. | 69 | 1/0 | 27.89 | Zhao et al., 2016. |
| AT5G05760 | SYP31, Qa-SNARE | **SYP31** and SYP32 play partially redundant roles in pollen development by modulating protein trafficking and Golgi structure. | 439 | 4/0 | 19.85 | Rui et al., 2021. |
| AT1G15880 | GOS11, Qb-SNARE | — | 1154 | 1/0 | 28.03 | — |
| AT2G45200 | GOS12, Qb-SNARE | **An** essential host factor for PD targeting of P3N-PIPO protein of potyvirus. | 427 | 4/0 | 25.89 | Song et al., 2016. |
| AT2G36900 | MEMB11, Qb-SNARE | **Involved** in anterograde protein trafficking at the ER-Golgi interface. | 240 | 4/0 | 20.8 | Marais et al., 2015. |
| AT3G58170 | BET11, Qc-SNARE | **BET11** and BET12 are required for fertility and pollen tube elongation. | 380 | 3/0 | 18.71 | Bolaños-Villegas et al., 2015. |
| AT4G14600 | SFT11, Qc-SNARE | — | 621 | 1/0 | 47.52 | — |
| AT1G29060 | SFT12, Qc-SNARE | **Plays** an important role in salt and osmotic stress responses and functions in the salt stress response via sequestration of Na<sup>+</sup> in vacuoles. | 147 | 1/0 | 72.5 | Tarte et al., 2015. |
| AT1G11890 | SEC22, R-SNARE | **Crucial** for male gametophyte development and cytoskeleton dynamics during plant development. | 654 | 1/0 | 46.48 | El-Kasmi et al., 2011; Guan et al., 2021. |
| AT5G22360 | VAMP714, R-SNARE | **Involved** in endocytic recycling to dynamically control the level and localization of PIN proteins. | 918 | 3/0 | 19.66 | Gu et al., 2021. |
| AT4G32410 | CESA1 | **An** important component of cellulose synthesis complex which synthesizes primary wall cellulose. Involved in pollen grain formation and pollen tube growth. | 220 | 1/0 | 33.02 | Chen et al., 2010; Mendu et al., 2011. |
| AT2G21770 | CESA9 | **Involved** in secondary cell wall biosynthesis. Important for pollen grain formation. | 792 | 5/0 | 21.75 | Chen et al., 2010; Stork et al., 2010. |
| AT5G52560 | USP | **A terminal** enzyme of the myo-inositol oxygenation pathway to UDP-glucuronic acid, a precursor for cell wall biosynthesis. | 1200 | 1/0 | 26.8 | Geserick and Tenhaken, 2013. |
| AT1G10290 | DRP2A | **DRP2A** and DRP2B function coordinately in post-Golgi pathways participating in transport of callose, a major component of the cell plates and PT cell wall. | 335 | 5/3 | 30.18 | Backues et al., 2010. |
| AT1G59610 | DRP2B | **Ditto.** | 335 | 7/4 | 30.18 | Ditto. |
| AT3G12160 | RABA4D | **Essential** for regulation of pollen tube tip growth in *Arabidopsis*. It is transported in exocytic vesicles to the apical tip of pollen tubes where it promotes tip growth | 262 | 2/0 | 19.8 | Szumliński and Nielsen, 2009. |
| AT3G51600 | LTP5 | **A small** secreted peptide from both pollen and the pistil, plays a role in pollen tube tip growth and in pistil function. It plays important roles in maintaining the cell polarity at the tube tip and adhesion-mediated guidance perhaps by interactions with pectins. | 347 | 5/0 | 29.04 | Chae et al., 2009. |

—, no reference. Abbreviations: Hit, peptide\_hit\_number; score, peptide\_score; BET11 (BS14a), bet1p/sft1p-like protein 14a; CESA1, cellulose synthase 1; DRP2A, dynamin related protein 2A; GOS11, Golgi SNARE 11; LTP5, Lipid transfer protein 5; MEMB11, membrin 11; SEC22, secretion 22; RABA4D, Ras-related brain GTPases A4D; SEC31B, secretory 31B; SYP31, syntaxin of plants 31; USP, UDP-sugar-pyrophosphorylase; VAMP714, vesicle-associated membrane protein 714.

In addition to the vesicle transport regulators, some intine/primary cell wall synthesis-related factors, CESA1, CESA9, UDP-sugar-pyrophosphorylase (USP), Dynamin related protein2A (DRP2A) and DRP2B, were also identified to associate with AtSYP32 (Table 3). As descripted above, CESA1/3/6 are involved in primary cell wall biosynthesis (Polko and Kieber, 2019). CESA2/5/9 and CESA6 are partially redundant (Desprez et al., 2007; Persson et al., 2007), and CESA10 has sequence homology with CESA1 (Griffiths et al., 2015) (Supplemental Table S2). USP is a terminal enzyme of myo-inositol oxygenation pathway to UDP-glucuronic acid, a precursor for cell wall biosynthesis (Geserick and Tenhaken, 2013); DRP2A and DRP2B function coordinately in post-Golgi pathways participating in transport of callose, a major component of the cell plates and PT cell wall (Backues et al., 2010) (Table 3). In addition to these factors, Fasciclin-like arabinogalactan protein3 (FLA3) specifically expressed in pollen grain and tube, is reported to be involved in pollen development and intine formation (Li et al., 2010) (Supplemental Table S2). RT-qPCR detection results indicate that apart from *DRP2A* and *DRP2B*, the expression levels of *CESA1*, *CESA2*, *CESA3*, *CESA9*, *CESA10*, *USP* and *FAL3* decreased significantly in *atsyp32+/-* mutants and *AtSYP32* RNAi lines compared with that in wild type (Figure 7C), suggesting that biosynthesis of intine/primary cell wall was disrupted.

Two other AtSYP32-associating proteins, RABA4D and Lipid transfer protein5 (LTP5), are the regulators of polar vesicle transport in pollen tube (Table 3). RABA4D GTPase, expressed specifically in pollen, is important for PT tip growth (Szumlanski and Nielsen, 2009). LTP5, a secretory peptide from both pollen and pistil, participates in PT tip growth (Chae et al., 2009). In addition to these two factors, RHO-related protein from plants1 (ROP1), Pollen receptor like kinase2 (PRK2), ROP1 enhancer1 (REN1) and EXO70C2 are reported to regulate polar transport in PTs (Supplemental Table S2). In *Arabidopsis*, ROP (Rho-like GTPases from plants) GTPases are key regulators of polar growth in PTs and other cells. ROP1 controls PT tip growth (Li et al., 2008); PRK2, a receptor-like protein kinase, regulates ROP1 signaling pathway via interacting with RopGEF1 and ROP1 (Chang et al., 2013); REN1, localized at PT apical PM and exocytic vesicles, deactivates ROP1 and maintains the apical ROP1 cap (Hwang et al., 2008); and EXO70C2 contributes to PT optimal tip growth (Synek et al., 2017). The expression levels of these genes reduced significantly in *atsyp32-1+/-* and/or *AtSYP32* RNAi lines compared with wild type (Figure 7D), further proving that AtSYP32-mediated vesicle transport contributes to PT tip growth.

Rupture of pollen grains and PTs (Figure 2E) as well as the significant alteration of expression of above genes (Figure 7, A-C) in *atsyp32+/-* mutants and *AtSYP32* RNAi lines indicate an essential role of AtSYP32 in integrity of pollen wall and PT cell wall. Therefore, we detected the expression of genes of the regulatory machineries. As introduced above, RALF4/19-LRX-AUN1 and RALF4/19-BUPS/ANX-MRI are two distinct pathways but converge to fine-tune PT CWI (Boisson-Dernier et al., 2015; Mecchia et al., 2017; Ge et al., 2017; Franck et al., 2018; Li and Yang, 2018; Wang et al., 2018; Ge et al., 2019). RT-qPCR determination indicated that the expression levels of *LRX11*, *RALF19*, *AUN1*, *BUPS1/2*, *ANX1/2* and *MRI* reduced significantly in *atsyp32+/-* mutants and *AtSYP32* RNAi lines compared with that in wild type (Figure 7E), while, that of *LRX10* and *RALF4* altered slightly but significantly in *atsyp32+/-* or *AtSYP32* RNAi lines, but *LRX8* and *LRX9* expression didn’t change obviously compared with that in wild type (Supplemental Figure S5A). These results suggest that AtSYP32 is required for maintenance of PT CWI probably via modulating RALF4/19-LRX-AUN1 and/or RALF4/19-ANX/BUPS-MRI regulatory machineries.

Since the pollen tube growth and targeting to the ovule were disturbed (Figure 2, G and H), we checked the pollen tube guidance machinery. PRK6, a pollen-specific receptor-like kinase, binds to the attractants AtLURE1s for pollen tube guidance. PRK6-AtLURE1-mediated signaling regulates micropylar pollen tube attraction (Zhong et al., 2019; Liu M. et al., 2021). XIUQIUs are also pollen tube attracting peptides (Zhong and Qu, 2019). The expression of *PRK6*, *XIUQIUs* and *LURE1s* decreased significantly in *atsyp32+/-* mutants and *AtSYP32* RNAi lines compared with that in wild type (Figure 7F), suggesting AtSYP32 may modulate PRK6-attractants signaling pathways. Taken together, AtSYP32 may regulate PT cell wall biosynthesis and PT CWI miantenance via modulating vesicle trafficking.

### AtSYP32 may coordinate with partners to regulate pollen wall development and maintenance of PT CWI

To determine AtSYP32 partners upon regulation of cell wall development and PT CWI, we firstly performed yeast two hybrid (Y2H) analysis to detect interactions between AtSYP32 and its functionally relevant factors related to pollen wall development and PT CWI maintenance (Figure 7). As expected, as a Golgi-localized Qa-SNARE, AtSYP32 physically interacted with Qb-SNAREs MEMB11/12 and GOS11/12, Qc-SNAREs BET11/12 and SFT11/12, and R-SNAREs VAMP714 and SEC22, respectively (Supplemental Figure S6A), suggesting that AtSYP32 may form different complex with these SNARE proteins to regulate vesicle trafficking in different occasions. AtSYP32 also interacted with the COPII coatomer SEC31B which is essential for pollen wall development by regulating secretion in tapetal cells (Supplemental Figure S6A), indicate that AtSYP32 might accept arriving COPII vesicle via interacting with COPII subunit SEC31B and R-SNARE SEC22. Beyond our predictions, AtSYP32 also physically interacted with the XyG xylosyltransferase XXT5, and PT CWI maintenance factors, RALF19, LRX11 and ANX2, respectively (Supplemental Figure S6B).

To confirm *in vivo* interactions between AtSYP32 and these factors, we performed a Split Luciferase Complementation Assay (SLCA) using tobacco leaves. The results indicate that apart from ANX2, AtSYP32 interacted with SEC22, SEC31B, BET12, LRX11, RALF19 and XXT5 in plant cells, respectively (Figure 8A). Since UGD2 showed self-activation in Y2H, we detected the interaction by SLCA but didn’t have positive result. To elucidate at which organelle AtSYP32 interacts with these factors, we performed Bimolecular fluorescence complementation (BiFC) assay upon tobacco leaves. The confocal images indicate that AtSYP32 interacted with SEC22, SEC31B, BET12, LRX11, RALF19 and XXT5 on the Golgi apparatus, respectively (Figure 8B-8J). These results further proved that AtSYP32 coordinates with these factors to regulate the pollen wall development and PT CWI.

**Figure 8.**
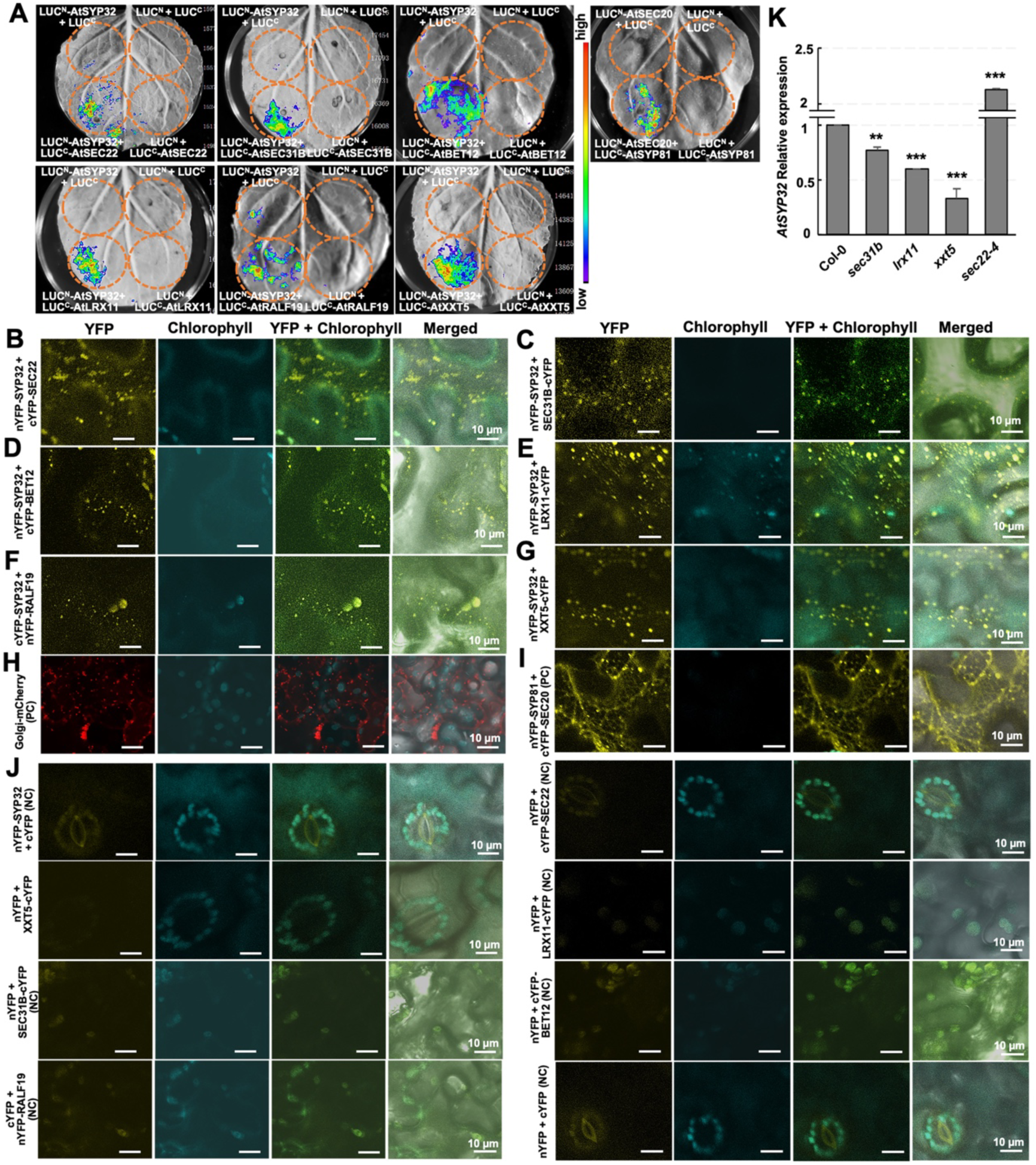
*In vivo* interaction of AtSYP32 with the partner proteins. A, SLCA analysis of AtSYP32 and the partner proteins. Plasmid combinations are as shown at the bottom-left of each panel. LUC^N^+LUC^C^, LUC^N^-AtSYP32+LUC^C^, LUC^N^+LUC^C^-interested proteins served as negative controls. The circles show the infiltration areas. B-J, BiFC analysis of AtSYP32 and its partners. Plasmid combinations are as shown. Positive and negative controls are as shown in (H-J). K, Statistics of RT-qPCR determination of relative expression levels of *AtSYP32* in Col-0, *sec31b, lrx11*, *xxt5* and *sec22-4*. Three independent experiments per sample, four technical replicates per experiment. **, *P* < 0.01; ***, *P* < 0.001; Student’s *t*-test. NC, negative control; PC, positive control.

To explore functional relevance between AtSYP32 and the partners in plants, we isolated *sec31b*, *lrx11*, *xxt5* and *sec22-4* mutants to detect *AtSYP32* expression. RT-qPCR determination indicate that *AtSYP32* expression reduced significantly in *sec31b*, *lrx11* and *xxt5*, while elevated dramatically in *sec22-4* compared with that in wild type (Figure 8K), suggesting tight functional relevance between *AtSYP32* and *SEC31B*, *LRX11*, *XXT5* or *SEC22*, respectively. We further detected the expression levels of the genes related to pollen wall biosynthesis and PT CWI maintenance in these mutants. The results indicate that the expression levels of most genes also altered significantly in *sec31b*, *lrx11*, *xxt5* and *sec22-4* compared with that in wild type, some gene expression were almost absent (Figure 7, right side in each panel), suggesting their functional relevance with each other.

Interacting factors regulate stability of each other. When one is defective, the stability of the other is usually affected. To validate the interaction of AtSYP32 with its partners in pollen tubes, we performed RNA probe labeling to detect transcription levels of *LRX11*, *XXT5* and *SEC31B* using pollen tubes collected from Col-0 and *atsyp32-1+/-* mutant. First of all, we confirmed that *AtSYP32* expression declined dramatically in *atsyp32-1+/-* (> 60%) compared with that in Col-0 (Figure 9, A and H), indicating that the experimental performance was reliable. As expected, the fluorescence intensities of *LRX11*, *XXT5* and *SEC31B* decreased significantly in *atsyp32-1+/-* pollen tubes compared with that in Col-0 (Figure 9, B-D, and H), indicating downregulation of *AtSYP32* affected the transcription of these genes. Since the Golgi integrity is disrupted in *syp31 syp32-1* double mutant (Rui et al., 2021), we detected the transcription level of a Golgi resident, *KATAMARI1* (*KAM1*) (Tamura et al., 2005). The result indicate that *KAM1* expression was reduced significantly in *atsyp32-1+/-* pollen tubes compared with that in Col-0 (Figure 9, E and H), suggesting downregulation of *AtSYP32* affected the Golgi functions. The same change occurred to a PM marker, *SYP132* (Ichikawa et al., 2014) (Figure 9, F and H), suggesting the secretory pathway was perturbed in *atsyp32-1+/-*. However, the transcription level of an ER resident, *SYP81* (Wang et al., 2022), didn’t alter obviously compared with that in Col-0 (Figure 9G, 9H), suggesting downregulation of *AtSYP32* didn’t affect the Golgi-to-ER retrograde transport severely. These results suggest that *AtSYP32* affects transcription of *LRX11*, *XXT5* and *SEC31B* in pollen tubes; meanwhile, *AtSYP32* modulates the Golgi stability and secretion efficiency.

**Figure 9.**
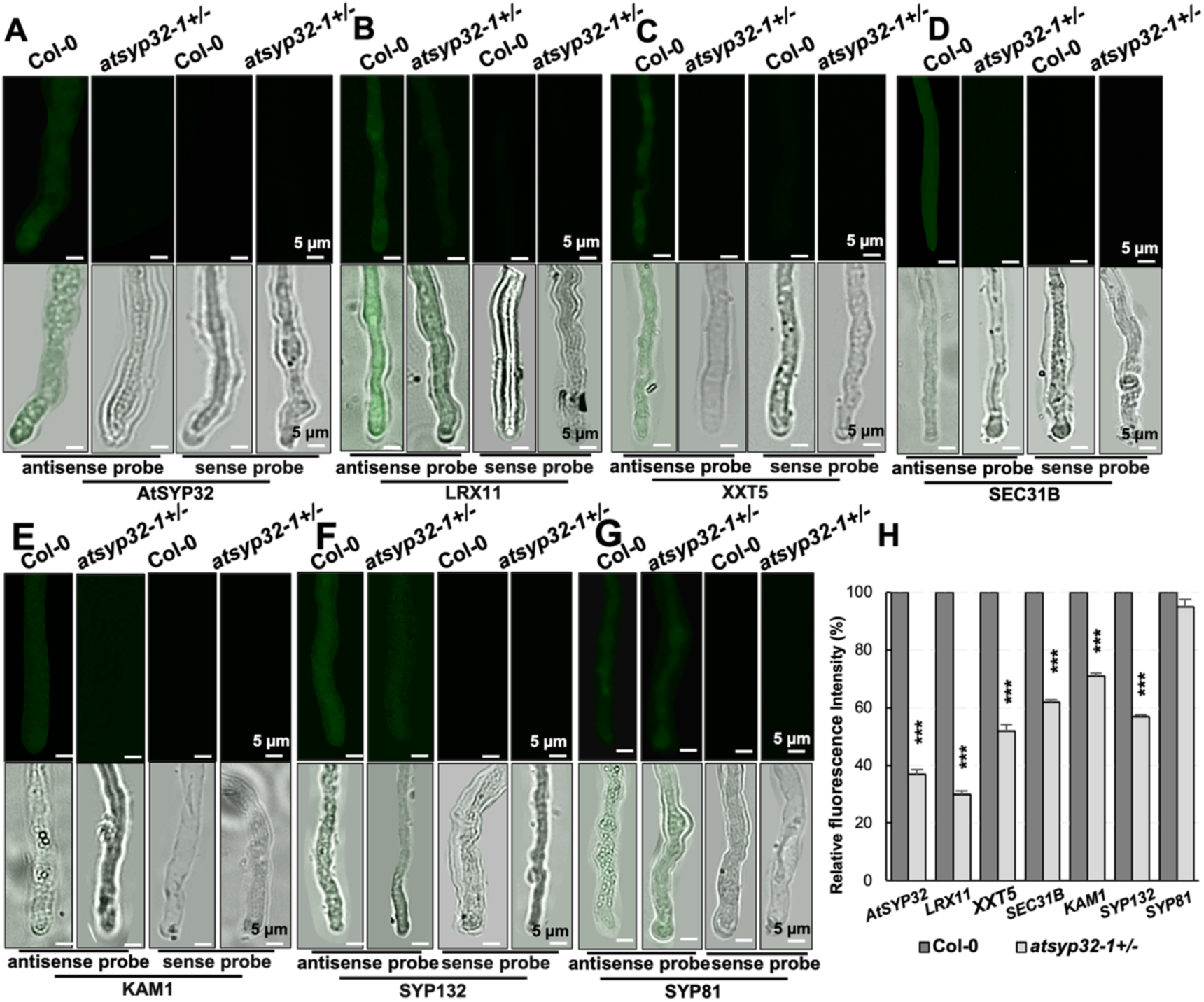
Detection of *in vivo* transcription level of *AtSYP32* and its partners in pollen tubes. A-G, RNA probe labeling detection of transcription levels of *AtSYP32* (A), *LRX11* (B), *XXT5* (C), *SEC31B* (D), *KAM1* (E), *SYP132* (F), and *SYP81* (G) in pollen tubes. H, Statistics of relative fluorescence intensities shown representatively in (A-G). Values are means ± SD (n ≥ 30), three biological replicates per sample. ***, *P* < 0.001; Student’s *t*-test.

## Discussion

The cis-Golgi syntaxin AtSYP3 family has two homologues, AtSYP31 and AtSYP32. Recent study demonstrate that AtSYP31 and AtSYP32 play partially redundant roles in pollen development by modulating intra-Golgi trafficking and Golgi morphology. In our study, we clarified that AtSYP32 plays an essential role in pollen wall development and PT CWI maintenance via regulating early secretory pathway and exocytosis of PT cell wall biosynthetic materials.

### AtSYP32 regulates early secretory pathway

AtSYP32 interacted with some vesicle trafficking regulators including SNAREs and COPII coatomer. Golgi-localized Qb-SNAREs MEMB11/12 and GOS11/12, Qc-SNAREs BET11/12 and SFT11/12, and R-SNAREs SEC22 and VAMP714, physically interacted with AtSYP32 (Supplemental Figure S6A), suggesting AtSYP32 may form distinct SNARE complexes to regulate anterograde transport from the ER to the Golgi. It is not clear under what conditions what combination of complex will be formed. As an important R-SNARE, SEC22 is demonstrated to be essential for the ER and Golgi integrity, may regulate anterograde transport and is crucial for pollen development (El-Kasmi et al., 2011; Guan et al., 2021), but the molecular mechanism underlying SEC22-regulated pollen development is elusive. Considering the co-expression and interaction with AtSYP32, SEC22 might be involved in pollen wall development, but it needs further investigation to prove the hypothesis.

Since anterograde and retrograde transport between the ER and the Golgi are interconnected processes, if one direction is blocked, as a consequence, another one is also affected. Therefore, the function disturbance of the ER and Golgi are usually coupled. The ER and Golgi morphology were seriously disturbed in *sec22-4*, a knockdown allele (Guan et al., 2021), suggesting that SEC22 is essential for anterograde and retrograde transport. The collapsed ER morphology in *atsyp32+/-* mutants and *AtSYP32* RNAi lines resembled that in *sec22-4* mutant, and downregulation of Golgi residents such as KAM1 and XXT5 suggest disordered Golgi functions. All these imply that AtSYP32 is required for both anterograde and retrograde transport between the ER and the Golgi, directly or indirectly. That SEC22 interacts with AtSYP32 rather than with AtSYP31 (Guan et al., 2021) (Supplemental Figure 6A) raises the question of what other subunits are in the SNARE complex that regulates the development of pollen wall and PT CWI. The Qc-SANRE BET11/12 are required for fertility and PT development (Bolaños-Villegas et al., 2015), and an *in vitro* studyt has suggested that BET11 and BET12 tend to form a distinct quaternary SNARE complex with different Golgi SNAREs (Tai and Banfield, 2001). Moreover, Qb-SNARE MEMB12 is identified as a partner of BET12 (Chung et al., 2018). The SLCA and BiFC analyses validated interaction of AtSYP32 with SEC22 or BET12 in plant cells (Figure 8). All these findings raise a potential quaternary SANRE complex composed of Qa-AtSYP32, Qc-BET12, R-SEC22, and probable Qb-MEMB12, which may regulate pollen wall development and PT CWI via modulating early secretory pathway (Supplemental Figure S8A). Identification of distinct AtSYP32-SNARE complex will help to uncover the biological significances of AtSYP32-mediated pathways.

SEC31B regulates pollen wall development via controlling early secretory pathway in tapetal cells (Zhao et al., 2016). The interaction between AtSYP32 and SEC31B may facilitate uncoating of the arriving COPII vesicles, and the trans-SNARE complex promotes COPII vesicles unloading cargoes which contain the important regulators of Golgi morphology and functions. Many kinds of proteins are synthesized in the ER and transported to the Golgi apparatus for post-translational modification. Then, some resident in the Golgi and some are transported to the vacuole, PM or apoplast. Therefore, the Golgi, functioning as a hub for vesicle transport pathways,

### AtSYP32 is required for secretion of biosynthetic materials and functional proteins of pollen wall and PT cell wall

A striking phenomenon in *atsyp32+/-* mutants, *AtSYP32* RNAi and *AtSYP3132* RNAi lines is the large amount of ectopic vesicles retained in the pollen cytoplasm (Figure 3, I and J; Supplemental Figure S3, B and C), indicating secretory pathway was seriously blocked. Among the ectopic vesicles, the probably ER-derived double membrane-bound vesicle, EXPO, has been demonstrated to fuse with the PM and release an exosome which contains cell wall biosynthetic enzymes and materials, such as a lignin methylase S-adenosylmethionine synthetase 2 (SAMS2), the arabinogalactan glycosyltransferases for AGP *O*-glycosylation, and proteoglycans (summarized by van de Meene et al., 2017). The glycosyltransferases (GTs) are localized at the ER or the Golgi apparatus, the ER-localized type initiate protein glycosylation, and the Golgi-localized type complete the glycosylation. For example, Galactosyl transferase 29A (GALT29A), GALT31A and Glucuronosyltransferase 14 (GlcAT14A) complete glycosylation of AGPs at the Golgi apparatus (summarized by van de Meene et al., 2017). It is illustrated that the GTs and AGPs are transited through the EXPO vesicles (Poulsen et al., 2014). EXPO-mediated transport of proteins are usually leaderless secretory proteins (LSPs), and the traffic route belongs to Unconventional protein secretion (UPS) (van de Meene et al., 2017), in other words, EXPO serves to secret proteins that cannot enter the conventional secretory pathway into the apoplast. Another potential UPS route is mediated by the specialized MVBs which are secretory compartments possibly fuse with the PM rather than being a prevacuolar compartment/late endosomes (Meyer et al., 2009; Ding et al., 2014). Callose has been detected in MVBs in epidermal cells attacked by pathogens (Mendgen, 1994), and the callose synthases Glucan synthase-like 5 (GSL5) is also found to be targeted to the PM via MVB-mediated UPS route (Jacobs et al., 2003; Nishimura et al., 2003; An et al., 2006).

Different from GSL5, GSL1 which is involved in pollen tube formation is delivered via the Conventional protein secretion (CPS) pathway (Brownfield et al., 2008). CPS route is responsible for secretion of proteins with an N-terminal leader sequence. These proteins are synthesized in the rough ER, modified in the ER and/or the Golgi and then exocytosed into the apoplast via SVs. SVs deliver cargoes from the TGN to the PM/apoplast, and different types of SVs contain different cargoes, including CESAs, GTs, xyloglucan and pectins (summarized by van de Meene et al., 2017). In plants, most of the polysaccharide synthases and GTs are localized at the Golgi or the PM in CPS route (Driouich et al., 2012; Oikawa et al., 2013; Wilson et al., 2015), highlighting the importance of CPS in biosynthesis and secretion of cell wall polysaccharides. The abnormal accumulation of numerous EXPOs, SVs, MVBs and other unclassified vesicles in the pollen cytoplasm in *atsyp32+/-* mutants and *AtSYP32* RNAi lines indicate that both CPS and UPS routes were severely disrupted due to AtSYP32 down-regulation. The significant decline of the intine thickness in the mutants and RNAi lines (Figure 3, G, H and K) reflected the effects of the blocked exocytosis. The significant decline of the cellulose contents (Figure 6G) and the expression levels of *CESAs*, *XXT5*, *UGD2*, *GlcAK*, *USP* and *FAL3* (Figure 7), the genes responsible for intine/primary wall biosynthesis, verified the disturbance on intine/primary wall deposition in the mutants and RNAi lines. Furthermore, the XXXG XyG and pectins delivered by SVs were accumulated inside the PTs (Figure 6, C and E) narrating the failed secretion. Excessive accumulation of XXXG XyG might feedback and inhibit the transcription of *XXT5*. The arabinogalactan glycosyltransferases responsible for AGP *O*-glycosylation are transported by EXPO (Poulsen et al., 2014). The traffic jam in *atsyp32-1+/-* PTs caused failure of the enzymes reaching the Golgi and resulted in incomplete glycosylation of AGPs, which may lead to degradation of AGPs. The abnormal retention of EXPOs in pollen cytoplasm and reduction of AGPs abundance in PTs of *atsyp32-1+/-* (Figure 3, H-J; 6D and G) support this possibility. All these results demonstrate that AtSYP32 is crucial for secretion of biosynthetic materials and functional proteins of pollen wall and PT cell wall.

### AtSYP32 is crucial for tapetum development

The *Arabidopsis* tapetum is of the secretory type, and plays an important role in the exine and pollen coat formation (Quilichini et al., 2015). The tapetosomes in tapetal cells are derived from the ER, and contain a large amount of lipidic components which will form the pollen coat (Shi et al., 2015). The collapse of ER morphology will lead to dysfunction of the ER, and subsequently affect tapetosome formation. The occurrence of abnormally large tapetosomes in *atsyp32+/-* mutants, *AtSYP32* RNAi and *AtSYP3132* RNAi lines may due to the transport failure of inhibitors for tapetosome fusion, such as oleosin proteins which play an essential role in tapetosome formation and protein relocation to the pollen coat (Lévesque-Lemay et al., 2016). Since synthesis of lipids is mainly in the ER (Shi et al., 2015; Zhao et al., 2016), dysfunction of the ER will also affect the inclusion of tapetosomes. Determination of the contents of the lipidic components will help to understand the importance of the ER in synthesis of these substances.

The elaioplasts in tapetum are derived from proplastids (Piffanelli et al., 1998). The reduced amount and ambiguous morphology of elaioplasts in *atsyp32+/-*, *AtSYP32* RNAi and *AtSYP3132* RNAi lines (Figure 4, A-C) suggest that AtSYP32-mediated vesicle trafficking might participate in the differentiation of elaioplasts from proplastids and vesicle trafficking in proplastids. In *sec31b* tapetal cells, elaioplast development is also retarded probably due to the postponed differentiation from proplastids to elaioplasts and reduced efficiency of COPII transport in proplastids (Zhao et al., 2016; Liu et al., 2021). These phenotypes in *atsyp32+/-* and *sec31b* mutants combined with the interaction between AtSYP32 and SEC31B illustrate that AtSYP32 may modulate the formation of elaioplasts. While, the hypothesis needs further experimental evidences validation.

### Functions of AtSYP31 and AtSYP32 have a division

AtSYP32 has a homologue protein AtSYP31 with 46% Identities and 62% Positives, but their functions seem to be quite different. In Y2H analysis, AtSYP31 didn’t interact with any of XXT5, RALF19, LRX11, ANX2, MEMB11, BET11 or SEC22, all of which interacted with AtSYP32; on the other hand, similar to AtSYP32, AtSYP31 interacted with SEC31B and the other SNAREs identified by pull down-LC-MS/MS (Supplemental Figure S6, A and B; Table 3), suggesting that AtSYP31 plays an inherent function as Golgi-syntaxin that mediating anterograde transport from the ER to the Golgi. While, different from *atsyp32*+/-mutants, *atsyp31* mutants didn’t have male sterility, since the seed fertility and the pollen viability were normal (Figure 1E, 2A; Supplemental Figure S1H). Moreover, that *syp31 syp32* mutations can be complemented by *pSYP32:SYP32* rather than by *pSYP32:SYP31* transgenes (Rui et al., 2021) evidenced their functional differences in pollen development.

The amino acid sequence comparison indicate that both proteins have the N-terminal motif (syntaxin 5N), SANRE domain and transmembrane domain (Supplemental Figure S7A). The SANRE domain and transmembrane domain in AtSYP32 and AtSYP31 are relatively conservative, but the N-terminal motifs have lower conservation (Supplemental Figure S7B). Moreover, AtSYP31 has a di-acidic motif essential for its ER export, Golgi targeting and an interaction with the COPII machinery (Melser et al., 2009), however, AtSYP32 doesn’t have this motif (Supplemental Figure S7B), suggesting their functional differences. Phylogenetic tree analysis indicates that AtSYP32 and AtSYP31 are in different branches (Supplemental Figure S7C). The cis-elements in the two promoters have some difference, e.g. the ‘endosperm expression’ element is in AtSYP32 promoter but is absent in AtSYP31 one; while, the ‘defense and stress responsive’, ‘circadian control’ and ‘auxin-responsive’ elements are in AtSYP31 promoter rather than in AtSYP32 one (Supplemental Figure S7D). These information indicate that during evolution, the regulatory patterns and functions of AtSYP32 and AtSYP31 had a division. On the other hand, many evidences proved that AtSYP31 and AtSYP32 are partially functional redundant in pollen development, since the phenotypes of *AtSYP3132* RNAi lines were more serious than those of *atsyp32+/-* mutants and *AtSYP32* RNAi lines (Figure 1, E and G; Figure 3, G-J; Supplemental Figure S2A, S3, A-D). AtSYP32 regulates cell wall development and PT CWI via directly binding to the important factors such as RALF19, LRX11, XXT5, SEC22 and BET12, which AtSYP31 never associate with (Figure 8; Supplemental Figure S6). However, both AtSYP31 and AtSYP32 bind to SEC31B, the essential regulator for pollen wall development, indicating AtSYP31 and AtSYP32 regulate an overlapped pathway and therefore showed partial function redundancy, but AtSYP31 plays a minor role.

### Biological significance of AtSYP32 interacting with PT CWI regulators

An unexpected result in this study was the interactions of AtSYP32 with XXT5, LRX11 or RALF19, respectively (Figure 8; Supplemental Figure S6B). XXT5 may act as a regulator or an organizer for the xyloglucan synthetic complex (Zabotina et al., 2008; Chou et al., 2012; Chou et al., 2015). XXT5 has a transmembrane domain, which enable it a relatively stable localization for efficiently organizing the complex. Therefore, we hypothesize that interaction of AtSYP32 with XXT5 might facilitate XXT5 efficient recruitment from the vesicles to the Golgi, and enriched at a certain region to organize the synthetic complex, thus promoting the synthesis, sorting and secretion of XyG (Supplemental Figure S8B). LRX11, a cell wall glycoprotein involved in maintenance of PT CWI, is localized in the cytoplasm and cell wall of PTs (Wang et al., 2018). The interaction between AtSYP32 and LRX11 on the Golgi apparatus (Figure 8E) raises a possibility that AtSYP32 may recruit LRX11 from the cytoplasm to the Golgi to promote LRX11 glycosylation, and subsequent sorting at the TGN for exocytosis (Supplemental Figure S8C). The secretory peptides RALF4 and RALF19 seem distributed in the cytoplasm and the cell wall (Ge et al., 2017). This localization pattern resembles that of LRX11, and the biological significance of AtSYP32-RALF19 interaction may conduct an efficient exocytosis of the peptide, similar to that of LRX11 (Supplemental Figure S8D). The loss of polar localization of highly methylesterified HGs and the pectins (Figure 6, A and E) and the significant alteration of expression of PT polar transport regulatory genes such as *LTP5*, *RABA4D*, *PRK2*, *REN1*, *EXO70C2* and *ROP1*, in *atsyp32+/-* mutants and *AtSYP32* RNAi lines (Figure 7D) suggest a crucial role of AtSYP32 in the polar transport and the polarity maintenance of the cell wall components.

In *atsyp32+/-* mutants, many PTs missed the micropyles (Figure 2G-I), and the expression levels of the pollen tube receptor gene *PRK6* and the maternal secreted attractant genes *LURE1s* and *XIUQIUs* decreased significantly (Figure 7F), suggesting the pollen tube guidance mechanism was disturbed. Moreover, the ratio of progenies from self-pollinated *atsyp32+/-* mutants exhibited no mutation : heterozygous : homozygous≈1:1:0, and that of cross-pollinated *atsyp32-1/-2+/-* mutants (♀) and wild type (♂) were 5:1 and 3:1, respectively (Table 1) indicate that sterility of *atsyp32+/-* mutants were from not only male gamete, but also maternal side, i.e. the effects of AtSYP32 dysfunction on plants was holistic. First of all, the secretion pathway in *atsyp32+/-* pollen tubes was blocked, such as exocytosis of AGPs and polysaccharides were disturbed severely (Figure 6). Then, the abnormal gene expression included not only PT CWI regulatory genes such as *RALF14*, *LRX11* and *PRK6* (Figure 7E, 7F, 9B), but also maternal attractant genes *AtLURE1s* and *XIUQIUs* (Figure 7F). We further detected the expression levels of *FER* and *RALF34*. FER is localized in the synergid cells, and RALF34 is a peptide ligand derived from female gametophyte (Ge et al., 2017). Both of them are essential for induction of pollen tube bursting and sperm release (Ge et al., 2017; Li and Yang, 2018). RT-PCR detection indicate that expression of *FER* reduced significantly in *atsyp32+/-* mutants, *AtSYP32* RNAi lines, *sec31b*, *lrx11*, *xxt5* and *sec22-4*, while that of *RALF34* decreased significantly only in *sec22-4* (Supplemental Figure S5B). These results indicate that the AtSYP32 function is not limited to pollen wall development, but also on the development of female gametes and plant somatic cells.

Due to the extremely poor fertility of the *atsyp32+/-* mutants, many attempts have been tried to cross the mutants with some markers of cell wall and vesicle transport, but no success has been achieved. Therefore, it is a pity that some occasions cannot provide data in protein levels.

In summary, our findings indicate that AtSYP32, a cis-Golgi-syntaxin, is essential for early secretion pathway and post-Golgi trafficking. AtSYP32 is localized in the tapetum, pollen grain and tube, regulating pollen wall development and maintenance of PT CWI via controlling secretory pathway. Furthermore, the interaction between AtSYP32 and XXT5 may promote the synthesis and secretion of XyG; and the interaction of AtSYP32 with LRX11 and RALF19 may promote their recruitment from the cytoplasm, modification at the Golgi and exocytosis to the apoplast.

## Methods and methods

### Plant materials and growth conditions

*Arabidopsis thaliana* ecotype Col-0 was used as wild-type plants. T-DNA-tagged lines were derived from Col-0. *atsyp32-1* (GABI_109A09), *atsyp32-2* (GABI_920F05) and *atsyp32-3* (SAIL_1293_A09), *atsyp31-1* (SALK_150783) and *atsyp31-2* (SALK_057421C) were obtained from the *Arabidopsis* Biological Resource Center (ABRC) at Ohio State University. *xxt5* (SALK_120831C) was obtained from the AraShare. *lrx11* (SALK_076356) (Wang et al., 2018), *sec31b* (SALK_103304) (Zhao et al., 2016) and *sec22-4* (SAIL_736_F03) (Guan et al., 2021) mutant lines were donated by corresponding groups. The seeds were surface-sterilized and sown either on soil or onto 0.8 or 1.2% agar with 1/2 Murashige and Skoog medium (PhytoTech, China) and 1% (w/v) sucrose. Plants were grown at 22°C under 16 h light/8 h dark photoperiod.

### RNA extraction, RT-qPCR and RT-PCR analysis

Total RNA was isolated using RNAiso Plus (9109, TAKARA, Japan). 0.5−1 μg of total RNA was reverse transcribed using the PrimeScript™ RT Master Mix (Perfect Real Time) (RR036A, TAKARA, Japan). RT-qPCR and RT-PCR was performed according to the manufacturer’s instructions. *ACT2* was used as an endogenous control for RT-qPCR and RT-PCT.

### Plasmid construction

An *AtSYP32* cds fragment was amplified from Col-0 using the *AtSYP32*-specific primers *AtSYP32 TOPO-F* and *AtSYP32 TOPO-R* and ligated into *pENTR/D-TOPO* vectors (Invitrogen, Carlsbad, CA, United States). To generate TAP-tagged (containing 9 x myc) *AtSYP32*-expressing transgenic plants, *the AtSYP32* cds fragment was transferred from the *AtSYP32* entry clone to the destination vector *pNaTAP* (Rubio et al., 2005) by an LR reaction (Invitrogen). For *generating AtSYP32* RNAi plants, a 151 bp fragment of *AtSYP32* cDNA was amplified using the primers *AtSYP32RNAi-F* and *AtSYP32RNAi-R*, and cloned into *the pENTR/D-TOPO* vector and subsequently subcloned into the destination vector *pK7GWIWG2* by the LR reaction. For *generating AtSYP3132* RNAi plants, a 252 bp fragment of *AtSYP32* cDNA was amplified using the primers *AtSYP3132RNAi-F* and *AtSYP3132RNAi-R*, and cloned into *the pENTR/D-TOPO* vector and subsequently subcloned into the destination vector *pK7GWIWG2* by the LR reaction. To generate *pAtSYP32:gAtSYP32* complementation plants, the primers *AtSYP32 comple-F1* and *AtSYP32 comple-R1* were used for promoter, and *AtSYP32 comple-F2* and *AtSYP32 comple-R2* were used for genomic sequence cloning, the fragments were cloned into *the pCAMBIA1301* vector by One Step Cloning Kit (Vazyme). For BiFC analysis, coding regions of *AtSYP32*, *BET12*, *RALF19*, *SEC31B*, *LRX11*, *XXT5*, *SEC22*, *SYP81* and *SEC20* were amplified and cloned into *pCAMBIA1300/35S-N-nYFP*, *pCAMBIA1300/35S-C-cYFP* or *pCAMBIA1300/35S-N-cYFP* vectors using One Step Cloning Kit (Vazyme). For SLCA, *pCAMBIA1300/N-Luc* and *pCAMBIA1300/C-Luc* vectors were used. The coding regions of above genes were cloned into vectors using One Step Cloning Kit (Vazyme).

The primers used here are listed in Supplemental Table S3.

### Generation of anti-AtSYP31 and anti-AtSYP32 antibodies

To prepare the antigen, the AtSYP31 cytosolic fragment corresponding to 1-750 amino acids was amplified using the primers *AtSYP31 TOPO-F* and *AtSYP31tm TOPO-R*, the AtSYP32 cytosolic fragment corresponding to 1-795 amino acids was amplified using the primers *AtSYP32 TOPO-F* and *AtSYP32tm TOPO-R*, ligated into *pENTR/D-TOPO* vectors, and subsequently introduced into the *pET32a* vector. Recombinant proteins were expressed in the *E. coli* BL21 strain, purified with a HiTrap chelating column, and entrusted to PhytoAB Inc., to generate polyclonal antibodies.

### Immunoblot analysis

Immunoblot analyses were performed as described previously (Li et al., 2006). Antibodies were diluted as follows: anti-AtSYP31, 1:1,000; anti-AtSYP32, 1:500; anti-actin (AS13 2640, Agriser), 1:1,500 and anti-myc (9E10:sc-40, Santa Cruz Biotechnology, Inc China. Shanghai), 1:2,000, respectively. The dilution of horseradish peroxidase-conjugated rabbit antibodies raised against rabbit IgG (ZB2301, ZSGB-BIO, China) was 1:5,000. Immunoreactive signals were detected using an enhanced chemiluminescence detection system (LAS−4000, FYJIFILM).

### Yeast two hybrid Assay

For yeast two-hybrid assay, the cds fragments of the interested genes (Supplemental Figure S5) were amplified and fused in-frame downstream of the GAL4 activation domain in the pGADT7 vector or downstream of the GAL4 DNA binding domain in the pGBKT7 vector. Positive control AtSYP81 and AtSEC20 constructs were generated in our previous study (Li et al., 2006). The paired constructs were introduced into strain AH109 of *S. cerevisiae* (Clontech) and selected on SD/−Leu/−Trp plates. The interactions were examined on SD/−Leu/−Trp/−His/−Ade plates.

### Pull down assay

Pull-down assays were performed as described previously (Li et al., 2013) using an iMACS epitope tag protein isolation kit (anti-c-myc, Miltenyi Biotec). Two grams of 10-days-old seedlings of *TAP-AtSYP32*-expressing (*AtSYP32 OE*) line were used. The binding beads were used for Shotgun liquid chromatography-tandem mass spectrometry (LC-MS/MS) analysis.

### Shotgun LC-MS/MS analysis

LC-MS/MS analysis was performed as described previously (Guan et al., 2021).

### AtSYP32 co-expressional gene analysis

The gene sets that are co-expressed with *AtSYP32* (AT3G24350) were identified using ATTED-II (https://atted.jp/). Specifically, the top 200 genes were extracted and the logit score (LS) was ranged from 9.3 to 4 (Obayashi et al., 2018).

### Gene and protein structural analyses

The AtSYP31 and AtSYP32 protein functional domains were identified and annotated in SMART (Simple Modular Architecture Research Tool) (http://smart.embl-heidelberg.de/), and pictures were drawn using TBtools software. DNAMAN8 was used to compare the amino acid sequences of AtSYP31 and AtSYP32. Parameter is the default parameter. At the same time, combined with protein structure analysis, Syntaxin-5N, SNARE domain and transmembrane domain were added in the comparison results. Search with SNARE domain (PF05739) in the pfam database (https://pfam.xfam.org/). A total of 12 SYP3 family proteins were identified in yeast (*Saccharomyces cerevisiae*), mouse (*Mus musculus*), human (*Homo sapiens*), maize (*Zea mays L.*), rice (*Oryza sativa L.*), alfalfa (*Medicago sativa L.*) and populus (*Populus trichocarpa Torr. & Gray*), and download the full protein sequence. Using MAGE11 software, using the maximum likelihood estimation method (Maximum Likelihood) and JTT + G mode, the number of tests is set to 1000, and other parameters are defaulted to construct the evolutionary tree. Then use the iTOL (https://itol.embl.de/) online website for beautification. *Arabidopsis* genome files and annotation files were downloaded from the Tair website (https://www.Arabidopsis.org), and 2,000 bp upstream of the start codons of *AtSYP31* and *AtSYP32* were extracted using TBtools for promoter analysis. Cis-acting regulatory elements were analyzed using PlantCARE (https://bioinformatics.psb.ugent.be/webtools/plantcare/html/), and finally imaged using TBtools.

### Scanning Electron Microscopy and Transmission Electron Microscopy analysis

Pollen grains were collected from freshly dehisced anthers and then mounted on scanning electron microscopy (SEM) stubs. The pollen grains were coated with palladium-gold in a sputter coater (JSM-7500F) and examined by SEM (JSM-7500F) at an acceleration voltage of 10 kV.

For ultrastructural observation, anthers containing mature pollen grains of stage 11-12 were fixed in 2.5% glutaraldehyde at 4°C, rinsed in 0.1 M phosphate-buffered saline (PBS, pH 6.8), and post-fixed in 1% OsO_4_ (dissolved in 0.1 M PBS). The anthers were embedded in Spurr’s resin for the cross-section procedure. Ultrathin sections (50-60 nm) were cut using a diamond knife on a Leica Ultracut ultramicrotome. Sections were double stained with saturated uranyl acetate and lead citrate and examined with a transmission electron microscope (H-7650; Hitachi).

### Pollen germination *in Vitro* and *in Vivo*

For pollen germination *in vitro*, mature pollen grains were spread on solid medium containing 0.01% H_3_BO_4_, 1 mM CaCl_2_, 1 mM Ca(NO_3_)_2_, 1 mM MgSO_4_ 10 % Sucrose and 0.5 % agarose. The samples were incubated for 4 h, and pollen germination results were examined with a microscope (Zeiss, AXIO Imager Z2).

For pollen germination *in vivo*, pollens from wild-type or the mutants were pollinated on wild-type stigma, respectively, after removing stamens. The pollinated pistils were collected at 48 h after pollination (hap) and stained by Aniline Blue.

### Immunofluorescence

After germination in liquid medium for 4 h, the pollen tubes were adhered to poly-L-lysine-covered glass slides, then fixed in 2.5% (w/v) polyformaldehyde in PIPES buffer [50 mM PIPES, 2 mM EGTA, 2 mM MgSO_4_, 5% (w/v) sucrose, pH 6.9] for 5 min, and then washed three times with PBS (0.2 M Na_2_HPO_4_, 0.2 M NaH_2_PO_4_, pH 6.9). The pollen tubes were incubated overnight at 4°C in dark with JIM5, JIM7, LM2 or LM15 antibodies (1:100), respectively. After wash five times with PBS, the pollen tubes were incubated for 3 h at 30℃ with DyLight 594 conjugated secondary antibody (1:100). The images were captured using a fluorescence microscope (Zeiss, AXIO Imager Z2).

### Cytochemical staining

For pollen and pollen tube cytochemical staining, the living samples were used. The pollen tubes were stained after germinating in liquid medium for 4 h. Alexander staining (Alexander, 1969) and DAPI staining (0.1 M sodium phosphate pH 7.0, 1 mM EDTA, 0.1% Triton X-100 [V/V], and 0.5 mg/mL DAPI) for pollen vitality, Aniline blue (0.1%, w/v) (Rui et al., 2021) for callose, and ruthenium red (0.01%, w/v) (Mecchia et al., 2017) for pectins, respectively. Images were captured with a fluorescence microscope (Zeiss, AXIO Imager Z2).

### Split Luciferase Complementation Assay (SLCA)

Overnight-cultured *A. tumefaciens* GV3101 (BC308-01, Biomed, China) harboring the constructs were resuspended (OD_600_ = 0.5) in infiltration buffer (10 mM MES pH 5.8, 10 mM MgCl_2_, and 100 μM acetosyringone) for 3 h in dark before infiltration. Equal volumes of the nLUC and cLUC suspensions were mixed and infiltrated into *N. benthamiana* leaves, which were placed in dark for 24 h and then transferred into the light for 48 h. Before fluorescence detection by a CCD camera (Vilber NEWTON7.0), the leaves were sprayed with 0.32 mg/mL D-Luciferin potassium salt in 0.1% Triton X-100 (Gold Biotechnology).

### Bimolecular Fluorescence Complementation (BiFC) Assay

*pCAMBIA1300/35S-N-nYFP*, *pCAMBIA1300/35S-C-cYFP* and *pCAMBIA1300/35S-N-cYFP* vectors with interested genes were used for transformation infiltration using tobacco leaves by *Agrobacterium tumefaciens* (GV3101). Leica laser scanning confocal microscope (SP8, Wetzlar, Germany) was used for fluorescence detection (YFP, excitation 514 nm, emission 518−582 nm).

### RNA probe labeling Assay

The probes of mRNA were synthesized as ∼300 bp transcripts, and then cloned into *pSPT18* vector. The T7/SP6 RNA polymerases were used to generate DIG-labeled probes by an *in vitro* transcription reaction using an NTP Labeling Mix (Roche, DIG RNA Labeling Kit SP6/T7, 11175025910). Pollen tubes adhered to poly-L-lysine-covered glass slides were fixed in 4% (w/v) polyformaldehyde for 30 min on ice, washed three times with PBS, incubated in 0.5% TritonX-100 / PBS solution for 10 min, washed three times. 70% precooled ethanol wash twice, and subsequently washed with 80%, 90% and 100% ethanol for dehydration. The prepared RNA probe [300 ng/μL, in hybridization solution (2 × SSC, 125 ng/μL salmon sperm DNA, 0.25% SDS, 10% Dextran Sulfate, 50% deionized formamide)] was denatured at 75℃ for 10 min. Added 70 μL hybridizing solution to the slide, and incubated at 37℃ overnight. Then, rinsed with 2×SSC/50% formamide solution at 37℃ for 5 min, with 2×SSC solution at 37℃ for 3 times, with 4×SSC/0.1% Tween-20 solution at RT for 5 min, finally used 4×SSC/4% BSA/0.1% Tween-20 sealing at 37℃ for 30 min. Then, incubated with rhodamine labeled antibody (Roche, 11207741910) (1:100) at 37℃ for 2 h. Images were captured with a fluorescence microscope (Zeiss, AXIO Imager Z2).

### Quantification of Cellulose

The first segments of main stems of 8-week-old plants were collected. Two grams of oven-dried stem powder was re-suspended in 80% (v/v) ethanol and heated at 50°C for 20 min. After centrifugation at 12,000 rpm for 10 min, the pellet was collected for quantification of cellulose using Cellulose Content Quantification Kit (Grace Bio., G0715W48, Suzhou, China). The protocol was followed the instructions.

### Accession Numbers

GenBank/EMBL accession numbers and *Arabidopsis* Genome Initiative locus identifiers for the genes mentioned in this article are as follows: *AtSYP32* (AT3G24350), *AtSYP31* (AT5G05760), *AtSYP81* (AT1G51740), *AtSEC20* (AT3G24315), *AtSEC22* (AT1G11890), *AtSEC31B* (AT3G63460), *AtLRX8* (AT4G08875), *AtLRX9* (AT1G49490), *AtLRX10* (AT2G15880), *AtLRX11* (AT4G33970), *AtXXT5* (AT1G74380), *AtGLcAK* (AT3G01640), *AtUGD2* (AT3G01640), *AtRABA1C* (AT5G45750), *AtRAB1A* (AT5G47200), *AtRABA4A* (AT5G65270), *AtRABA4B* (AT4G39990), *AtRABA4D* (AT3G12160), *AtVAMP721* (AT1G04750), *AtVPS45* (AT1G77140), *AtVPS46* (AT1G17730), *AtDL1C* (AT1G14830), *AtGOS11* (AT1G15880), *AtGOS12* (AT2G45200), *AtMEMB11* (AT2G36900), *AtMEMB12* (AT5G50440), *AtBET11* (AT3G58170), *AtBET12* (AT4G14455), *AtSFT11* (AT4G14600), *AtSFT12* (AT1G29060), *AtVAMP714* (AT5G22360), *AtCESA1* (AT4G32410), *AtCESA2* (AT4G32410), *AtCESA3* (AT5G05170), *AtCESA9* (AT2G21770), *AtCESA10* (AT2G25540), *AtUSP* (AT5G52560), *AtDRP2A* (AT1G10290), *AtDRP2B* (AT1G59610), *AtLTP5* (AT3G51600), *AtFLA3* (AT2G24450), *AtPRK2* (AT2G07040), *AtPRK6* (AT5G20690), *AtREN1* (AT1G77570), *AtEXO70C2* (AT5G13990), *AtROP1* (AT3G51300), *AtXIUQIU1* (AT5G50423), *AtXIUQIU2* (AT5G18403), *AtXIUQIU3* (AT5G18407), *AtXIUQIU4* (AT5G48605), *AtLURE1.1* (AT5G43285), *AtLURE1.2* (AT5G43510), *AtLURE1.3* (AT5G43513), *AtLURE1.7* (AT4G08869), *AtLURE1.8* (AT4G08875), *AtANX1* (AT3G04690), *AtANX2* (AT5G28680), *AtBUPS1* (AT4G39110), *AtBUPS2* (AT2G21480), *AtMRI* (AT2G41970), *AtAUN1* (AT3G05580), *AtFER* (AT3G51550), *AtRALF4* (AT1G28270), *AtRALF19* (AT2G33775), *AtRALF34* (AT5G67070).

## Supplemental data

The following materials are available in the online version of this article.

**Supplemental Figure S1.** Phenotypic analysis of *atsyp32+/-* and *atsyp31* mutants, *AtSYP32* RNAi and *AtSYP3132* RNAi lines.

**Supplemental Figure S2.** Knockdown of *AtSYP32* affected pollen vitality.

**Supplemental Figure S3.** Ultrastructural observation of pollen grains.

**Supplemental Figure S4.** Observation of callose deposition.

**Supplemental Figure S5.** The relative expression of the PT cell wall-related genes.

**Supplemental Figure S6.** Yeast two hybrid analysis of AtSYP32-interacting factors.

**Supplemental Figure S7.** Phylogenetic and structure analysis of SYP31 and SYP32 homologs.

**Supplemental Figure S8.** AtSYP32 function diagram.

**Supplemental Table S1.** Information of *atsyp32+/-*, *atsyp31*, *AtSYP32* RNAi and *AtSYP3132* RNAi lines.

**Supplemental Table S2.** Information of the genes related to PT cell wall biosynthesis and integrity maintenance.

**Supplemental Table S3.** Primers used in this study.

## Funding

This work was supported by the National Natural Science Foundation of China (32170279 and 31570246) and Fundamental Research Funds for the Central Universities (2572019CT03).

## Acknowledgments

We thank Prof. Rui Li (College of Life Science, Hebei Normal University) for providing seeds of *sec31b* and *lrx11* lines; Prof. Ikuko Hara-Nishimura (Kyoto University) for the gifts of *NaTAP* and *pK7GWIWG2* vectors; and Prof. Huazhong Shi (Depertment of Chemistry and biochemistry, Texas Tech University) for providing seeds of mCherry-AtSYP32 line.

## Author Contributions

L.L. conceived and supervised the project. L.L. and Y.L. designed the experiments. Y.L., X.Z., GT.Q., X.M., M.S., and K.G. prepared experimental materials and conducted experiments. GC.Q. performed the LC MS/MS and analyzed the data. Y.L. and X.M. analyzed the data. S.S. and M.W. conducted bioinformatics analysis. L.L. and Y.L. drafted the manuscript. J-K.Z., L.J. and W.S coordinated the project, and revised the manuscript. All authors contributed to manuscript revision, read, and approved the submitted version. The authors declare no competing interests.

